# Extracellular proteases are an essential public good supporting *Bacillus subtilis* growth through exogenous protein degradation

**DOI:** 10.1101/2023.02.08.527645

**Authors:** Thibault Rosazza, Lukas Eigentler, Chris Earl, Fordyce Davidson, Nicola Stanley-Wall

## Abstract

Bacteria encounter polymeric nutrient sources that need to be processed to support growth. *Bacillus subtilis* is a bacterium known for its adaptability and resilience within the rhizosphere and broader soil environment. Here we explore the role that a suite of extracellular proteases plays in supporting growth of *B. subtilis* when an extracellular heterologous protein (BSA) provides an abundant, but polymeric, food source. We confirm the essential role of extracellular proteases in this context and note the influence of the polymeric nutrient concentration on the yield of growth, but not on the relative level of extracellular proteases. We demonstrate the collective action of the extracellular proteases in supporting *B. subtilis* growth and evidence their use as a shared public good. Furthermore, we show that *B. subtilis* is subjected to a public good dilemma, but only in the context of using a polymeric food source. Using mathematical simulations, we uncover that this dilemma is driven by the *relative* cost of producing the public good. Collectively, our findings reveal how *B. subtilis* can survive in environments that vary significantly in terms of immediate nutrient accessibility. This information should inform steps to improve its efficacy as a biofertilizer in agricultural settings.

## Introduction

In the natural environment, there is the potential for bacteria to encounter a wide range of environmental conditions that differ in terms of nutrient availability and accessibility levels (1, 2). In contrast, laboratory experiments often use standardised levels of readily accessible nutrients that frequently do not reflect the physiology of the natural environments in which bacteria species reside (3). Soil habitats are one such example of an environment that can be associated with demanding growth conditions in which nutrient diversity, abundance, and accessibility vary (4). Several species of soil-associated bacteria are promising candidates for sustainable alternatives to fertilizers used in commercial agriculture (5). Therefore, understanding bacterial growth dynamics in these types of demanding environmental conditions will advance their development and efficacy.

One mechanism that microbes use to process complex, polymeric nutrient sources is the secretion of enzymes. The suite of enzymes required is linked to the ecological niche occupied by the producing strain (6-8). For example, *Vibrio cholerae* produces chitinase to break down chitin into oligosaccharides, which supports growth (9). Another example is the *Trichoderma reesei* cellulases, which are involved in depolymerizing plant cell wall polysaccharides to access carbon (10). In addition to the diversity of secreted enzymes that are encoded across different species, it has been shown that individual cells within bacterial communities can be metabolically heterogeneous (11). For example, spatially resolved single-cell transcriptomics has revealed that in *Pseudomonas aeruginosa* extracellular enzymes are produced in subpopulations of cells within an isogenic community (12). Furthermore, as these enzymes are secreted products, they are prone to exploitation by non-producers (cheaters) and are considered to be a “public good” (13). Social dynamics can lead to the occurrence of a *public good dilemma*; exploitation of the public good by non-producers leads to an increase in their relative density and consequently to a reduction in public good abundance, and eventually to population collapse (14).

*Bacillus subtilis* is a Gram-positive bacterium that colonises plant roots (15) and is well-known for its plant growth-enhancing properties (16). *B. subtilis* has been isolated from a broad range of environmental conditions, including many different soil environments, and the isolates exhibits high phenotypic variability (17). This variability reveals an extensive ability to adapt, which is in part due to its capability to grow using a broad range of carbon and nitrogen sources (18, 19). However, little is known about how this bacterium grows efficiently under environmental conditions that do not allow direct metabolization of nutrients. Despite this gap in knowledge, *B. subtilis* is currently commercialised and extensively used as a biofertilizer (20).

In this study, we focused on a group of eight extracellular proteases produced by *B. subtilis* (21). Six of the extracellular enzymes are within the serine-peptidases family (AprE, Bpr, Epr, Mpr, Vpr and WprA), while two are associated with a metallo-proteases family (NprB and NprE) (22). For many of these extracellular enzymes, a specific biological function has either been identified or has been hypothesised (23). The enzymes also mediate the extracellular degradation of proteins (23) and have been postulated to be a public good required to support growth (24). To the best of our knowledge, these conjectures remain unproven. Hence, we hypothesised that *B. subtilis* extracellular proteases would play a vital role in facilitating access to complex food sources that are contained in exogenous polymeric proteins. We compared NCIB 3610, a producer of all eight extracellular proteases (WT), and an otherwise isogenic mutant that lacks all eight genes responsible for extracellular protease production (Δ8). We determined the ability of extracellular proteases to degrade an exogenous protein, bovine serum albumin (BSA), and use it as a source of carbon and nitrogen to support growth. Furthermore, we determined that *B. subtilis* did not modulate extracellular protease production in response to changing levels of BSA in the growth medium or even when BSA was replaced by an accessible nutrient. We confirmed the *public good* property of this family of enzymes and by combining mathematical modelling with experimental approaches we predicted the impact of sharing this resource within a bacterial community containing differing proportions of extracellular enzyme producing and non-producing cells co-cultured in selected nutrient contexts.

## Materials and methods

### Strain construction

Strains used in this study were derived from *B. subtilis* isolate NCIB 3610 or NCIB 3610 *comI*^*Q12L*^ (stocked here as NRS6017) (25) (Table 1). To prepare competent cells, we used a media containing 60 mM K_2_HPO_4_, 37 mM KH_2_PO_4_, 95 mM D-Glucose, 3 mM sodium citrate dihydrate, 10 mM L-glutamic acid monopotassium salt, 0.1% (w/v) casein enzymatic hydrolysate, 3 mm MgSO_4_, 0.8 mM FeCl_3_ (25). A single colony was grown in 2 mL at 37°C, 200 rpm for 4.5 h. Then, 400 μL of this culture was mixed with 20 μL of gDNA (ranging from 100 ng/μL to 1 μg/μL) and incubated for 1.5 h at 37°C, 200 rpm. 100-200 μL of the sample was plated onto an LB plate (1% (w/v) tryptone, 0.5% (w/v) yeast extract, 1% (w/v) NaCl and select agar 1.5% (w/v)) with antibiotic for selection. Extracellular protease monoproducer strains were constructed by the insertion of an erythromycin resistance gene proximal to the wild type coding region in NRS6017, followed by the transfer of the allele to the Δ8 genome using the antibiotic selection marker. The introduction of the single extracellular protease coding region in the native location on the genome was assessed by PCR. See Table 2 for all primer sequences used in this study and Table 3 for all plasmids used in this study.

**Table 1.**
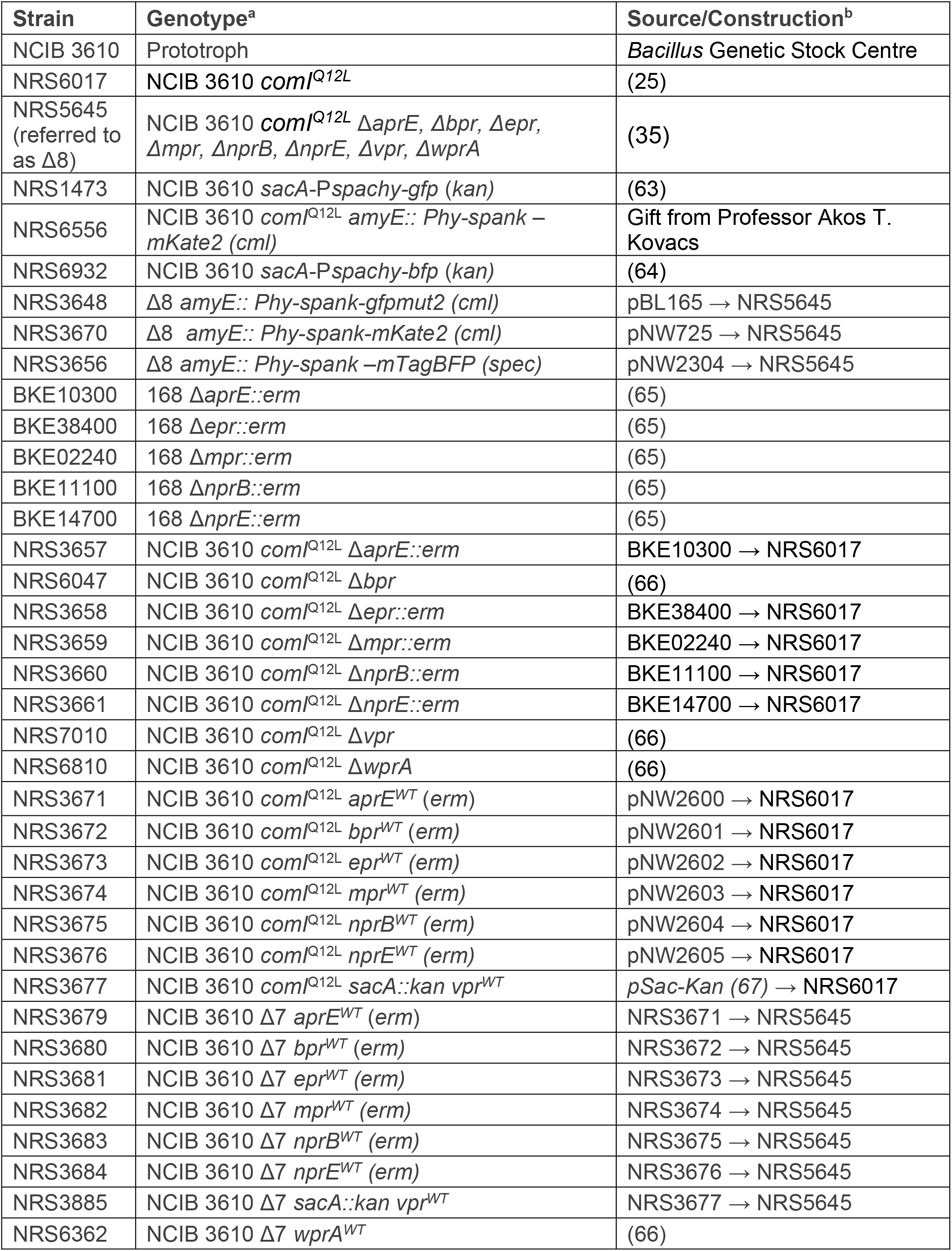
Strains used in this study. ^a.^ Drug resistance cassettes are as follows: *kan*, kanamycin resistance; *cml*, chloramphenicol resistance; *spec*, spectinomycin resistance; and *erm*, erythromycin resistance. ^b.^ Strain construction is denoted as DNA from donor strain transformed into recipient strain following the direction of the arrow (⟶). The reference is provided if the strain has previously been described. pNW and pDR numbers refer to plasmids (see Table 3) and BKE numbers refer to strains obtained from the BKE library (65).

**Table 2.**
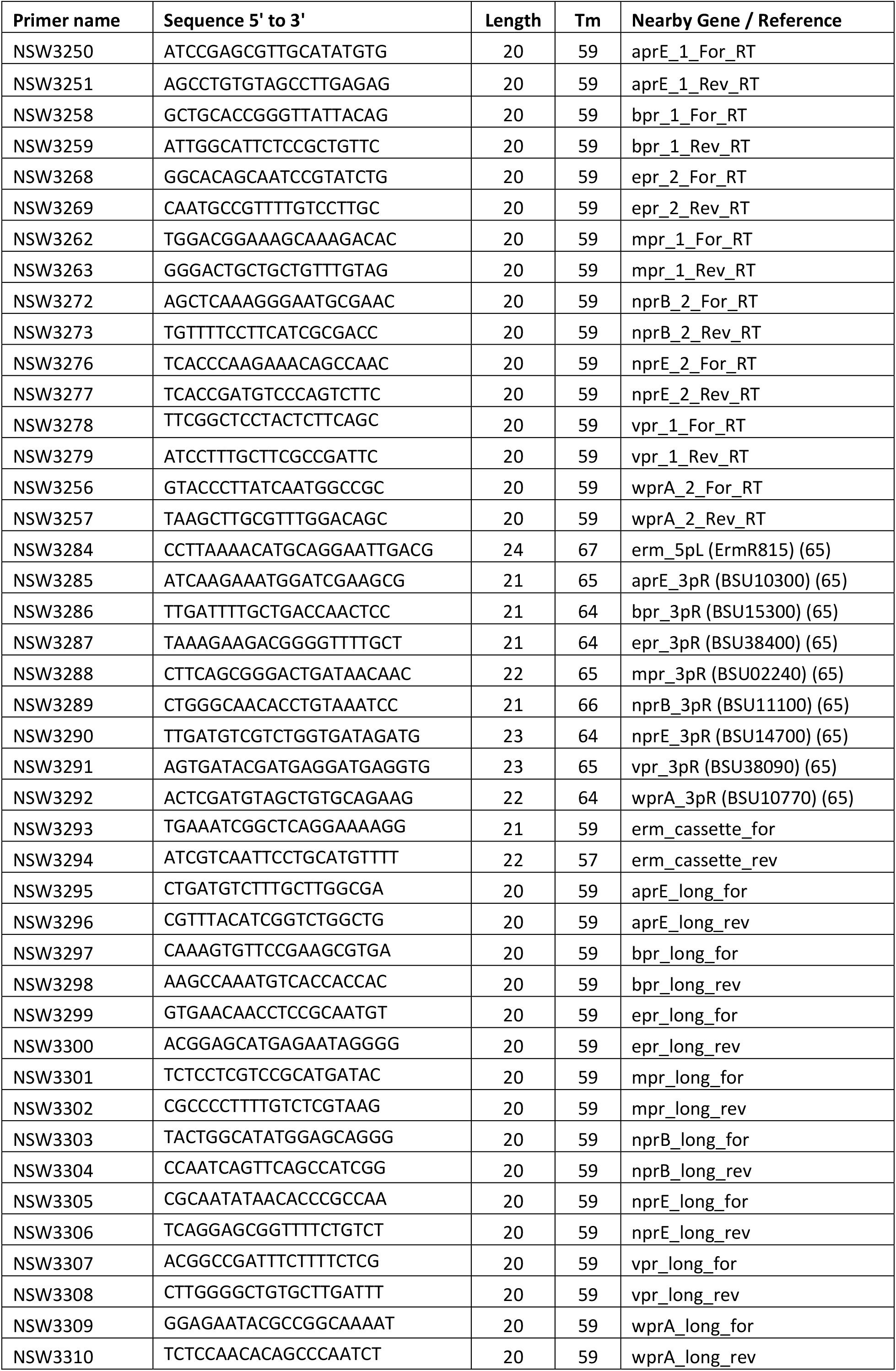
List of primers used in this study.

**Table 3.** List of plasmids used in this study. Synthetic cassettes were designed using NCIB 3610 genomic DNA as a template and erythromycin (*erm*) resistance gene (Supplementary data). pSac-Kan plasmid was used for integration of the kanamycin resistance gene at the *sacA* gene.

| Plasmid name | Background vector | Insert Fragment / Reference |
| --- | --- | --- |
| pBL165 | pBL165 | *Pspac-hy gfpmut2* (68) |
| pNW725 | pBL165 | *PtapA mKate2* |
| pNW2304 | pDR111 | *Phy-spank bfp* (64) |
| pNW2600 | pUC57 | *aprE-erm* cassette |
| pNW2601 | pCC1 | *bpr-erm* cassette |
| pNW2602 | pUC57 | *epr-erm* cassette |
| pNW2603 | pUC57 | *mpr-erm* cassette |
| pNW2604 | pUC57 | *nprB-erm* cassette |
| pNW2605 | pUC57 | *nprE-erm* cassette |
| pSac-Kan | pSac | *sacA-KanR* (67) |

### Genome Sequencing and analysis

Genome sequencing was provided by MicrobesNG (http://www.microbesng.uk). For sample preparation, single colonies of each strain to be sequenced were re-suspended in sterile PBS buffer (137 mM NaCl, 2.7 mM KCl, 10 mM Na_2_HPO_4_, 1.8 mM KH_2_PO_4_ pH 7.4) and streaked onto LB agar plates. The plates were incubated at 37 °C overnight and the following day. For short read sequencing, genomic DNA was extracted using QIAGEN kit (69504 – QIAGEN) and resuspended in EB buffer. For enhanced genome sequencing, the cells were harvested, placed into the barcoded bead tubes provided and sent to the MicrobesNG facilities. There, for each sample, three beads were washed with extraction buffer containing lysozyme and RNase A, incubated for 25 min at 37°C. Proteinase K and RNaseA were added and incubated for 5 min at 65°C. Genomic DNA was purified using an equal volume of SPRI beads and resuspended in EB buffer. DNA was quantified in triplicate with the Quantit dsDNA HS assay in an Eppendorf AF2200 plate reader. Genomic DNA libraries were prepared using Nextera XT Library Prep Kit (Illumina, San Diego, USA) following the manufacturer’s protocol with the following modifications: two nanograms of DNA instead of one were used as input, and PCR elongation time was increased to 1 min from 30 s. DNA quantification and library preparation were carried out on a Hamilton Microlab STAR automated liquid handling system. Pooled libraries were quantified using the Kapa Biosystems Library Quantification Kit for Illumina on a Roche light cycler 96 qPCR machine. Libraries were sequenced on the Illumina HiSeq using a 250 bp paired end protocol. Reads were adapter trimmed using Trimmomatic 0.30 with a sliding window quality cutoff of Q15 (26). De novo assembly was performed on samples using SPAdes version 3.7 (27), and contigs were annotated using Prokka 1.11 (28). Annotated draft assemblies of the sequencing results were acquired, whole genome sequencing data were visualised in Artemis software (29) and mutation predictions were determined using Breseq (30).

### Preparing cells for culture and assessing growth

Material from a -80°C glycerol stock of the required strains was streaked onto an LB plate and incubated O/N at 30°C. A single colony was used to inoculate 5 mL of LB and incubated O/N at 200 rpm at 37°C. 100 μL of the O/N culture was added to 5 mL of LB and incubated at 200 rpm at 37°C. After 4 h, the culture was centrifuged for 10 min at 4500 rpm. The cell pellet was resuspended using 1 mL of base MS media (5 mM K_2_HPO_4_, 5 mM KH_2_PO_4_, 100 mM MOPS pH 7.0) and OD_600_ was measured. The culture density was normalised to an OD_600_ of 1 by adding base MS media as required. In 50 mL Corning® tubes, 5 mL of base MS media was supplemented with metal mix (2 mM MgCl_2_, 700 μM CaCl_2_, 50 μM MnCl_2_, 50 μM FeCl_3_, 1 μM ZnCl2, 2 μM thiamine) and inoculated to an OD_600_ of 0.01. The growth media also contained 0.5% (w/v) glutamic acid, 0.5% (v/v) glycerol, or BSA at between 0.05-2% (w/v) as required. Note that the growth medium containing glycerol and glutamic acid is referred to as MSgg and has been used for a wide array of biofilm studies (31). Here we use it as a defined growth medium. The samples were incubated at 200 rpm at 37°C. The OD_600_ of the cultures was measured after 12, 24, 48, 72, or 96h incubation as indicated. An aliquot of the culture was sampled for spore or protease activity quantification. A separate culture tube was used for each time point.

### Measuring the percentage of spores

At 6, 12, 24, and 48h the cultures were collected and diluted at a 1/10 ratio in 1x PBS. Serial dilutions were performed and 100 μL was plated onto LB agar plates. This sample provided the total CFU in the culture. To specifically measure the fraction of spores, the serially diluted samples were heat-treated for 20 min at 80°C followed by 20 minutes at room temperature. 100 μL of the sample was plated onto LB plates after mixing thoroughly. The agar plates were incubated O/N at 37°C and the number of CFUs on each plate was enumerated to calculate the CFU/ml and CFU spores/ml.

### Extracellular proteases activity quantification

To measure the level of extracellular proteases a 1 mL sample from a planktonic culture was centrifuged for 10 min at 10,000 rpm. The supernatant was recovered and used with the protease fluorescent detection kit (PF0100 – Sigma-Aldrich). The protocol was adjusted from the manufacturers’ instructions in the following ways: 1) the samples were allowed to incubate for 4h, and 2) 20 μL of the trichloroacetic acid precipitation supernatant was used instead of 2 μL. The fluorescent signal at 485 nm was acquired using a 96 black well plate (Corning -CLS3603-48EA) and a Pherastar FSX plate reader (BMG Labtech) (protocol: endpoint fluorescence intensity, 485nm, 20 flashes, gain 100). The fluorescent signal was normalised relative to the OD_600_ of the culture at 48h.

### BSA digestion assay

A 1 mL culture supernatant from a 24h planktonic culture grown in media with glycerol and glutamic acid was centrifuged for 10 min at 10,000 rpm. For each sample, 500 μL of the supernatant was heat-treated 20 min at 80°C to ablate the enzymatic activity. For each condition, 15 μL of the culture supernatant (plus and minus heat treatment) was mixed with 20 μg of BSA contained in 15 μL (1.33 mg/mL stock solution). These samples were incubated at 37°C with shaking at 200 rpm for 12h. 10 μL of SDS loading dye (0.5 M Tris-HCl pH 6.8, 0.1 M EDTA, 15.5% (w/v) SDS, 3% (v/v) glycerol, 5% (v/v) β-mercaptoethanol, 1 mg bromophenol blue) was added to the samples which were heated for 5 min at 99°C. The integrity of the BSA was assessed after separation by 10% (w/v) SDS-PAGE and staining using Coomassie blue (ISB1L – Sigma-Aldrich). The resultant gel was imaged using an Azure 600 scanner (Azure Biosystems).

### Transwell^®^ assay

Cultures were prepared and normalised to an OD_600_ of 1 as described above. To physically separate the cell types, we used a Transwell® with a 0.4 μM pore size (10147291 – Fisher Scientific). For each strain combination, 1 mL of cell culture was added to the larger outer well and 250 μL was added to the small upper Transwell®. The samples were incubated at 30°C with no shaking in a plastic box (20cm by 10 cm by 10cm) filled with approx. 100 mL of water to keep the hygrometry level constant while not submerging the plates inside the box. After 3 days (glutamic acid and glycerol) or 9 days (BSA and glycerol) of incubation, both reflected light and fluorescence signals (485 nm and 620 nm) were captured using a Leica M165C stereoscope. Using FIJI software, a circular area encompassing the Transwell® and three circular areas within the outer well were drawn and fluorescence intensity signals were quantified.

### Co-culture of producers and non-producers

Cell cultures were prepared and normalised to an OD_600_ of 1 as described above. In a volume of 1 mL of MS base two strains were mixed in a ratio of 0:100, 25:75, 50:50, 75:25, 85:15, 95:5, and 100:0 (NRS1473:NRS3656). For each condition, 5 mL of media containing glycerol and BSA was inoculated at OD_600_ of 0.01 and incubated at 200 rpm at 37°C. At 6, 12, 24, and 48h the OD_600_ was measured, and a sample of each dilution was serially diluted in 1xPBS. 100 μl of the dilutions were plated onto LB plates and to distinguish the two strains the plates were imaged using an Azure 600 scanner (Azure Biosystems) to capture GFP and BFP fluorescent signals and link the colonies to each strain.

### Modelling the role of extracellular proteases

We used a continuum approach to describe the growth dynamics of bacterial cells, nutrient processing, and extracellular proteases production in well-shaken liquid cultures. Hence, we used a system of ordinary differential equations (ODEs) describing the interactions between the variables, which represent time-dependent densities: the extracellular proteases producing strain *W*(*t*) and its spores *W*_*s*_(*t*) that represent the wild type NCIB 3610; an extracellular proteases non-producing cheater strain *C*(*t*), and its spores *C*_*s*_(*t*) that represent the Δ8 mutant; an accessible nutrient *A*(*t*) that represents glutamic acid; a complex nutrient source *B*(*t*) that represents BSA; an intermediate nutrient *B*_*d*_(*t*) that represents the degraded BSA; and extracellular proteases *E*(*t*). Model simulations were run for times 0 ≤ *t ≤ t*_*end*_, where *t*_*end*_ represents the endpoint of our experimental assay. The system of equations used was

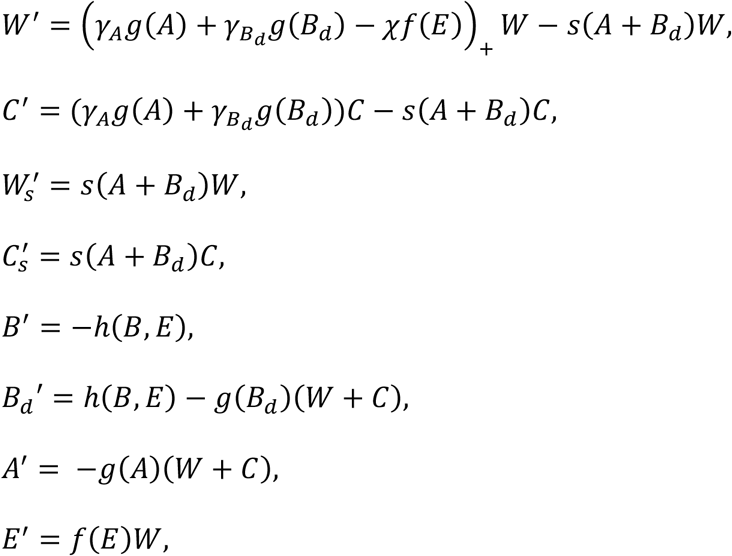

where

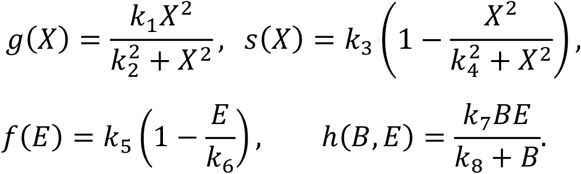

Both the extracellular enzyme producer *W* and non-producer *C* were assumed to grow in response to both the accessible nutrient source *A* and intermediate nutrient *B*_*d*_. The functions *g*(*A*) and *g*(*B*_*d*_) describe the nutrient consumption of each nutrient, respectively, assumed to be saturating and modelled using a Hill equation with Hill coefficient 2, maximum growth rate *k*_*1*_ > 0 and half saturation constant *k*_*2*_ > 0.

Corresponding growth functions (*γ*_*A*_*g*(*A*) and 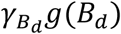), respectively) were assumed to differ only through the yield coefficients 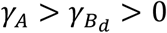. Each unit of the producer *W* was assumed to produce *f*(*E*) units of extracellular proteases per unit time. Due to negative feedback (the extracellular proteases degrade the quorum sensing signal ComX, which promotes extracellular protease production (32)), the production rate *f*(*E*) was assumed to be self-limiting and decrease from its maximum *k*_5_ > 0 to zero as the density *E* increased from zero to the carrying capacity *k*_6_ > 0. The penalty on growth borne by the producer was assumed proportional to the protease production rate and given by the term −*χf*(*E*)*W*. The parameter *χ* ≤ 0 was defined to be the *absolute cost* for extracellular protease production (a measure of the metabolic cost associated with producing one unit of extracellular protease per unit producer). We noted that in some of our experiments detailed below, cells entered a late death phase, due to nutrient exhaustion. We did not consider this in the model as it did not impact on the main objective, which was to uncover what determined the growth dynamics when nutrients were present (but potentially unavailable). Hence, we assumed the growth term in the first equation is non-negative. By definition, the non-producer was assumed to be free of growth penalty associated with extracellular protease production. Both producers and non-producers were assumed to sporulate at rate *s*(*A* + *B*_*d*_) in response to the total density of nutrients facilitating growth, *A* + *B*_*d*_. The sporulation rate was assumed to decrease from its maximum *k*_3_ > 0 to zero as the total nutrient density increased from zero following a Hill-type function with Hill coefficient 2 and half saturation constant *k*_4_ > 0. Finally, the conversion of the inaccessible base nutrient *B* to the intermediate nutrient *B*_*d*_ induced by extracellular proteases was assumed to follow Michaelis-Menten dynamics with maximum conversion rate *k*_7_ > 0 and half-saturation constant *k*_8_ > 0. The initial conditions for all cases were taken to be *W*(0) = *W*_O_, *C*(0) = *C*_O_, *W*_*s*_(0) = 0, *C*_*s*_(0) = 0, *B*_*d*_(0) = 0, *E*(0) = 0. Initial conditions for *B* and *A* differed across model scenarios. We represented growth media containing glutamic acid as the main nitrogen and carbon source by *B*(0) = 0, *A*(0) = *A*_*O,MSbg*_ > 0, and growth media containing BSA as the main nitrogen source by *B*(0) = *B*_*O,MSbg*_ > 0, *A*(0) = *A*_*O,MSbg*_ > 0, where *A*_*O,MSbg*_ was assumed to be much smaller than *A*_*O,MSgg*_. We highlight that in this second case (growth media containing BSA as the only nitrogen source) the initial amount of accessible nutrient *A* was small, but non-zero. This was to capture the observation that cells in the experimental assay displayed initial, fast growth that we concluded to be due to carryover of nutrients between the different growth conditions.

The model was numerically solved using Matlab’s ODE solver *ode15s*. Parameter values used are shown in Table 4. We were interested in qualitative agreement between the model and experimental assays and therefore did not estimate parameters using experimental data. However, we nevertheless ensured good qualitative fit across different nutrient conditions (see results). For visualisations, we normalised computed cell-densities using the density obtained for the in-silico WT in stationary phase strain in the growth medium representing glutamic acid.

**Table 4.**
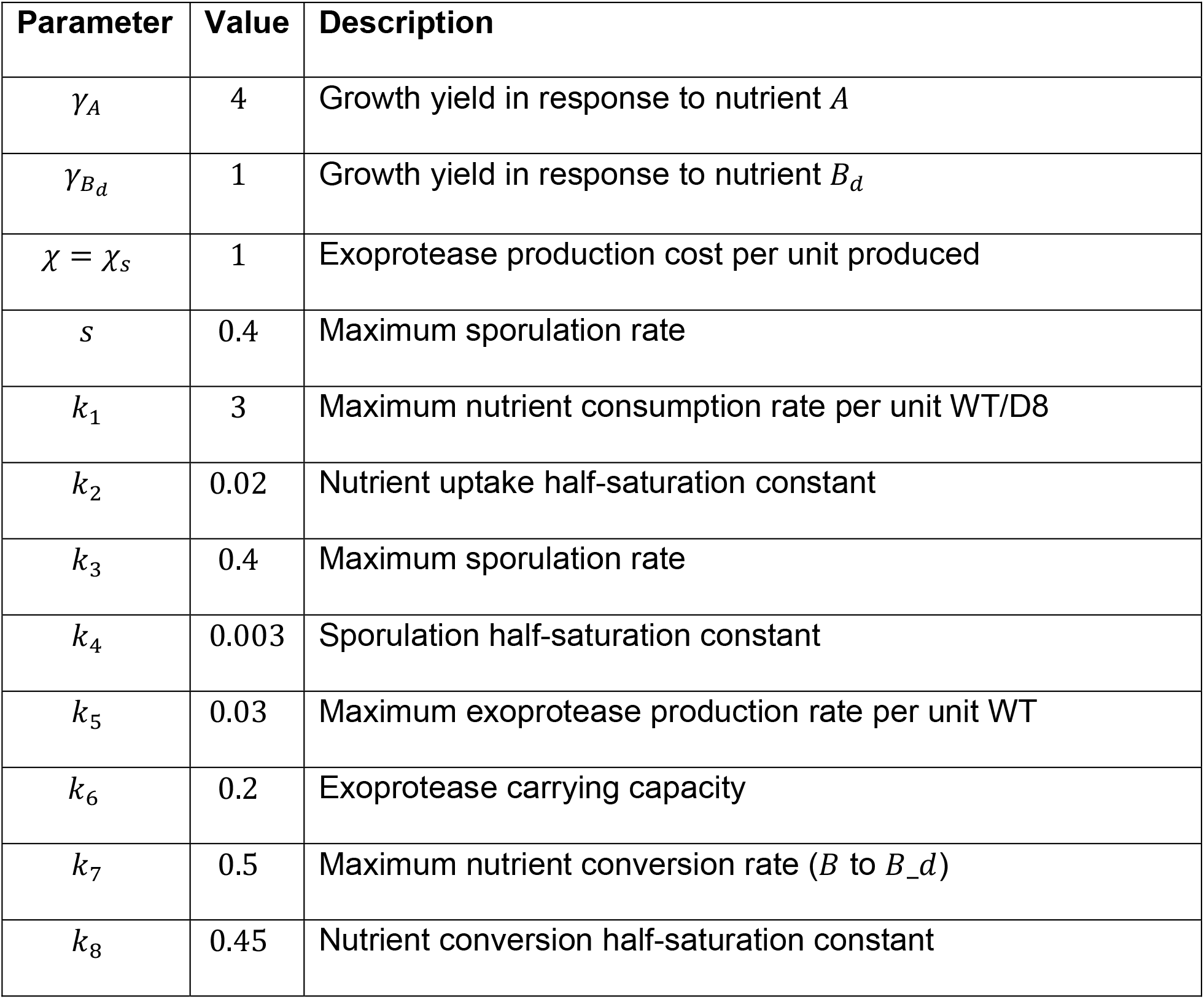
Model parameter values. Parameter values used (unless otherwise stated) in all simulations of our mathematical model are shown.

### The relative cost of extracellular protease production

We calculated the *relative cost* of extracellular protease production in our model as follows. First, we determined the time interval *T* in which the growth rate of *W*(*t*) was positive, i.e.,

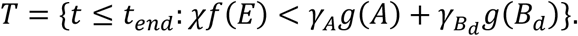

This represents the period over which wild type cells actively divide. We then defined the *total penalty* per unit of *W* of extracellular protease production during this time interval to be

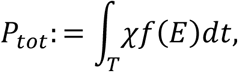

and the *total growth* per unit *W* in the absence of extracellular protease production to be

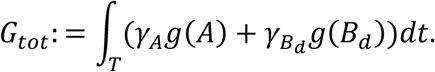

We then defined the *relative cost* of extracellular protease production per unit *W* during growth as the ratio

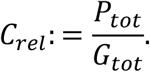

It is clear from the definition of *T* that the relative cost 0 ≤ *C*_*rel*_ ≤ 1.

### Statistical and data analysis

For group comparison, ANOVA test was performed. For mean comparison over multiple conditions, Tukey’s HSD test was performed. Data were analysed using Python 3.9 through Jupyter Notebook. All figures generated were generated using Matplotlib and Seaborn packages.

## Results

### *B. subtilis* extracellular proteases can digest the heterologous protein BSA

After growth of the wild type strain (WT, NCIB 3610) to stationary phase, we found that the suite of extracellular proteases contained in the culture supernatant (Fig. 1A) digested the heterologous bovine serum albumin (BSA) into oligomeric form (Fig. 1B). The observed BSA digestion was not present after heat-induced enzyme inactivation (Fig. 1B). We also showed that the culture supernatant derived from strain NRS5645, which lacks the coding regions for *aprE, bpr, epr, mpr nprB, nprE, vpr* and *wprA* (hereafter referred to as the “Δ8” strain), did not contain substantial proteolytic activity and did not cause any visible BSA digestion (Fig. 1A, 1B). We concluded that BSA would serve as a suitable polymeric nutrient source to assess growth dynamics.

**Figure 1.**
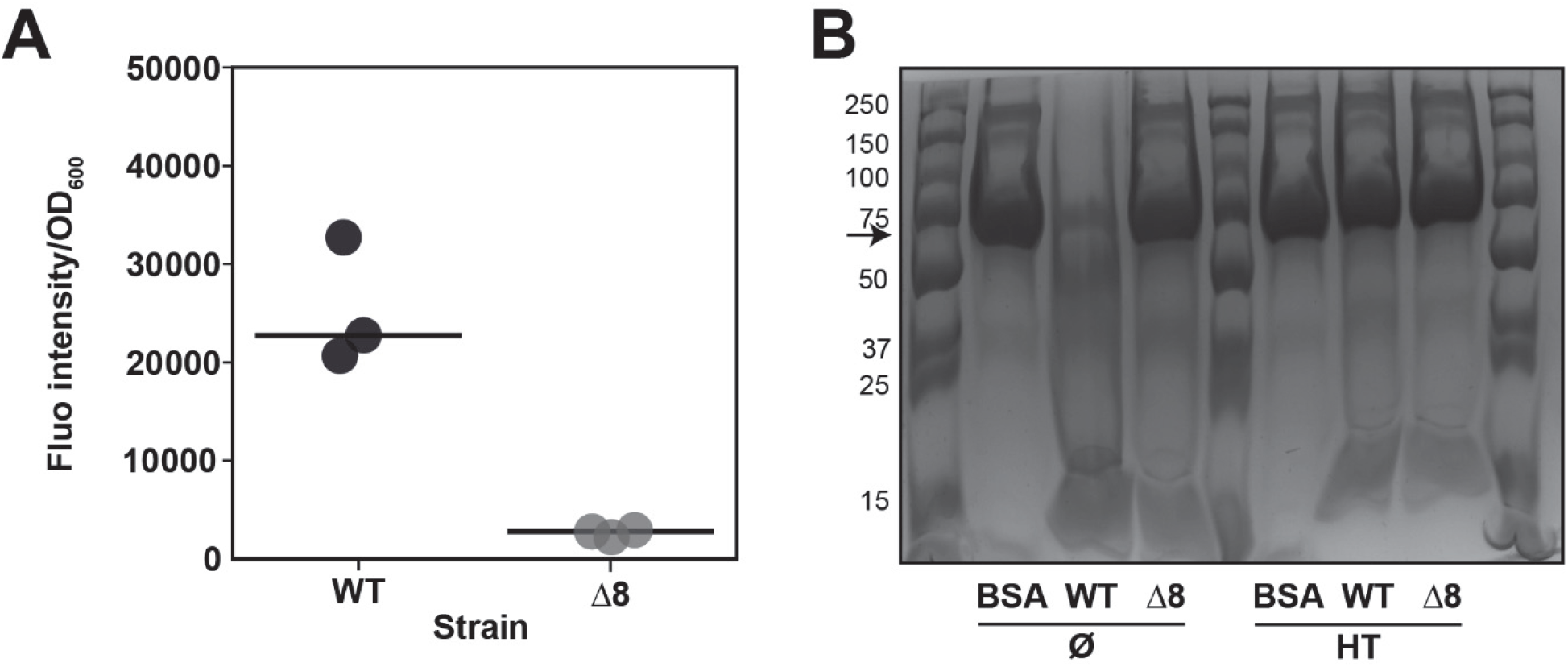
Extracellular proteases can digest BSA. **A**. Extracellular protease activity in glutamic acid 0.5% (w/v) and 0.5% glycerol (v/v) liquid monoculture culture of NCIB 3610 (WT) and NRS5645 (Δ8) strains, normalised to the OD_600_ of the culture when the supernatants were collected. Points represent fluorescence intensity/OD_600_ (n=3), and lines represent median. **B**. BSA digestion assay using the culture supernatant after monoculture culture of NCIB 3610 (WT) and NRS5645 (Δ8) strains before (Ø) and after heat-treatment (HT). BSA protein was mixed in water (BSA) or with culture supernatants for 12 hours at 37°C. Black arrow represents BSA molecular weight (69 kDa). A representative image of three independent experiments is shown.

### Extracellular proteases are required for growth when nutrients are in polymeric form

We monitored growth of the WT and Δ8 strains using four shaking culture conditions in which we varied the components containing nitrogen and carbon to modify the “complexity” of nutrient accessibility. We additionally quantified the proportion of spores in the cultures. We first compared the growth of the WT and Δ8 strains in a medium containing glutamic acid (a carbon and nitrogen source) and glycerol (a carbon source) that are both readily accessible nutrients (Fig. 2A, Fig. S1A). This is a medium that has been extensively used for growth of *B. subtilis* and is frequently referred to as “MSgg” (31). Each strain exhibited a period of rapid exponential growth that was followed by a short stationary phase and subsequently a death phase. For both strains, there was a low level of spores in the total population during the exponential growth phase and a high proportion of spores that formed during the stationary phase (Fig. 2B).

**Figure 2.**
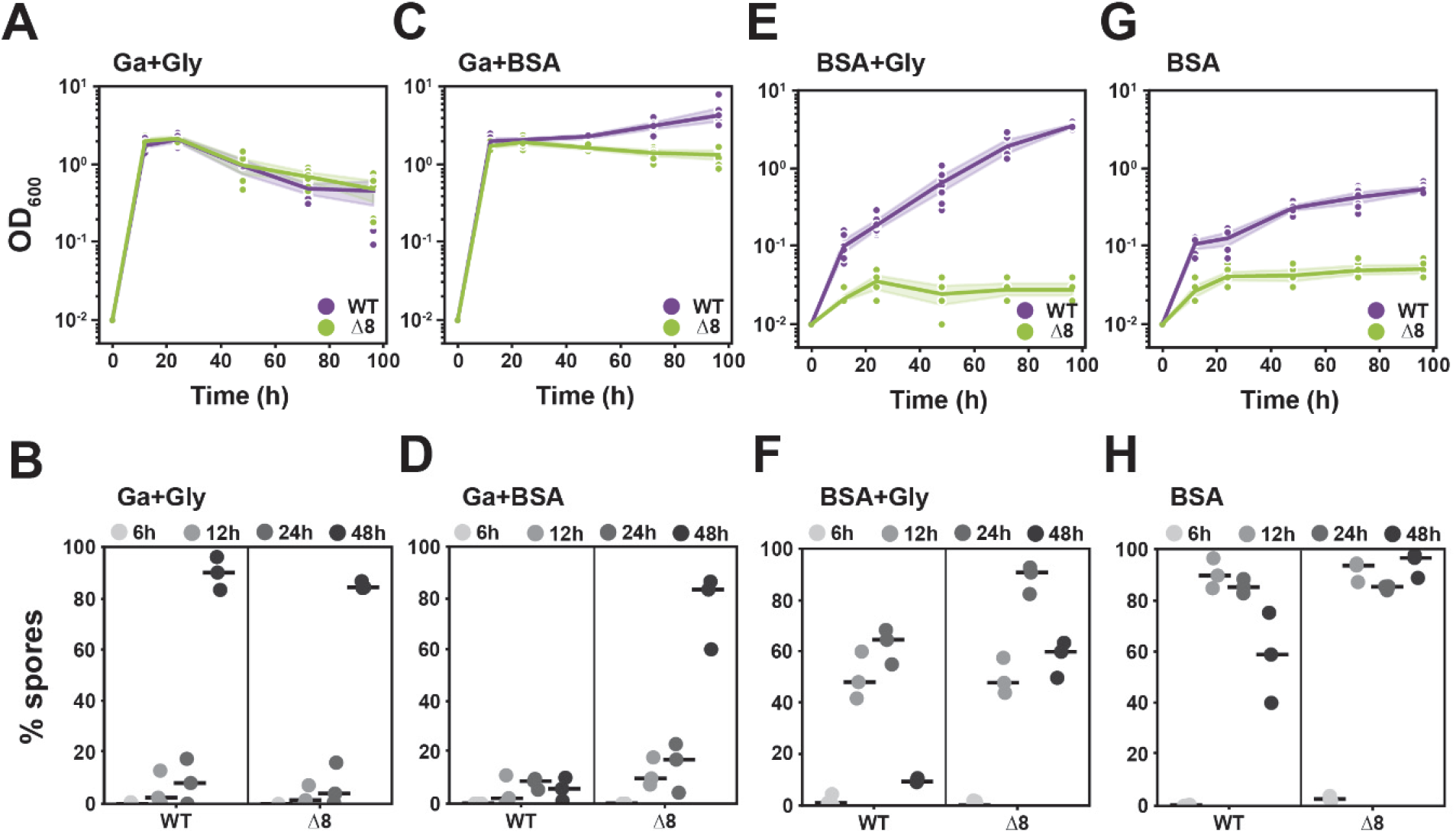
Differences in growth and sporulation with different nutrient accessibilities. **A, C, E, G**. Growth of NCIB 3610 (WT, purple) and NRS5645 (Δ8, green) in glutamic acid 0.5% (w/v) and 0.5% glycerol (v/v) (Ga+Gly) (A), 0.5% glutamic acid (w/v) and 1% BSA (w/v) (Ga+BSA) (C), 1% BSA (w/v) and 0.5% glycerol (v/v) (BSA+Gly) (E) and 1% BSA (w/v) (BSA) (G). Points represent OD_600_ values (n=3 biological replicates with 2 technical replicates), lines represent median and coloured areas represent CI 95%. **B, D, F, H**. Proportion of spores for each strain at different time points in monoculture grown in 0.5% glutamic acid (w/v) and 0.5% glycerol (v/v) (Ga+Gly) (B), 0.5% glutamic acid (w/v) and 1% BSA (w/v) (Ga+BSA) (D), 1% BSA (w/v) and 0.5% glycerol (v/v) (BSA+Gly) (F) and 1% BSA (w/v) (BSA) (H). Points represent % spores (n=3 biological replicates) and lines represent median.

When the growth medium was modified to contain glutamic acid and BSA (which are both sources of both carbon and nitrogen) (Fig. 2C), each strain displayed comparable periods of rapid exponential growth. However, after 48 hours of incubation, a difference in growth was observed. The extracellular protease-producing WT strain initiated a second, slower phase of sustained growth, while the extracellular protease-non-producing strain did not grow further. Moreover, the WT strain retained a low proportion of spores at all time points, while a significant proportion of the Δ8 population sporulated after the initial period of rapid growth (Fig. 2D). Note, when the WT strain was grown in culture medium that contained only glutamic acid, the total yield was lower than when BSA was also present (Fig. S1B). These findings allow us to conclude that the initial rapid growth exhibited by both strains is due to the cells using glutamic acid as the primary source of carbon and nitrogen. Moreover, our findings revealed that the sustained growth (displayed only by the WT strain) was facilitated by the presence of BSA, which was used as a secondary source of nutrients in an extracellular protease-dependent manner.

We next explored growth and sporulation in a growth medium that contained glycerol and BSA (Fig. 2E, Fig. S1C). Firstly, we observed that only the WT strain was able to sustain growth. Secondly, we observed that the doubling time of the WT strain was longer when compared to growth in glutamic acid and glycerol nutrient conditions (compare Fig. S1A and S1C). However, after five days the WT strain reached a higher yield compared to that attained in the glutamic acid and glycerol nutrient conditions (Fig. S1D). Thirdly, a significant proportion of the population of both the WT and Δ8 strains formed heat-resistant spores by 12 hours, but the level of spores reduced in the WT strain during later time points. This was consistent with an increase in growth (Fig. 2F).

Finally, when BSA was the sole source of carbon and nitrogen in the growth medium once more, only sustained growth of the WT strain was observed (Fig. 2G). When presented with this nutrient source, the WT attained a lower overall yield at the end of the experiment and a significant proportion of its population was in spore form. The Δ8 strain was not able to sustain growth and much of the population rapidly sporulated and remained in that state for the duration of the experiment (Fig. 2H). Collectively, our results prove the long-held conjecture that the extracellular proteases have a role in feeding.

### Two-phase growth response to increasing BSA level is mediated by saturating extracellular protease activity

We next explored if there was an impact on growth and extracellular protease production by the WT strain when the BSA concentration varied over a range of 0.05% to 2% (w/v). We used 0.5% (v/v) glycerol as an additional carbon source in all cases to enhance growth. Analysis of the exponential growth phase across each BSA concentration (Fig. 3A) revealed a saturation effect with broadly comparable growth rates after the BSA concentration exceeded 0.25% (w/v) (Fig. 3B). Below this threshold, the rate of growth decreased with decreasing BSA concentration (Fig. 3B). We compared the level of extracellular protease activity in the spent culture supernatant of the WT strain after growth at 48 hours. In all cases, the extracellular protease activity levels were comparable after normalisation to the yield (Fig. 3C). It is important to note that the extracellular proteases are stable in the culture supernatant for at least 24 hours (Fig S2A). Therefore, the values we measured represent the pool of extracellular proteases that had accumulated over time. As a control, we ensured that the presence of BSA in the growth medium did not interfere with the extracellular protease quantification process (Fig. S2B). Collectively, our data identify a two-phase response when *B. subtilis* is grown using a polymeric nutrient source. When nutrients levels are low, it is these that limit growth, as there are proteases available to degrade polymeric nutrients in the medium. Thus, increasing the level of nutrients results in an increase in exponential phase growth rate. However, when nutrients are abundant, then it is the extracellular proteases that are the limiting factor as further addition of nutrients does not increase extracellular protease production.

**Figure 3.**
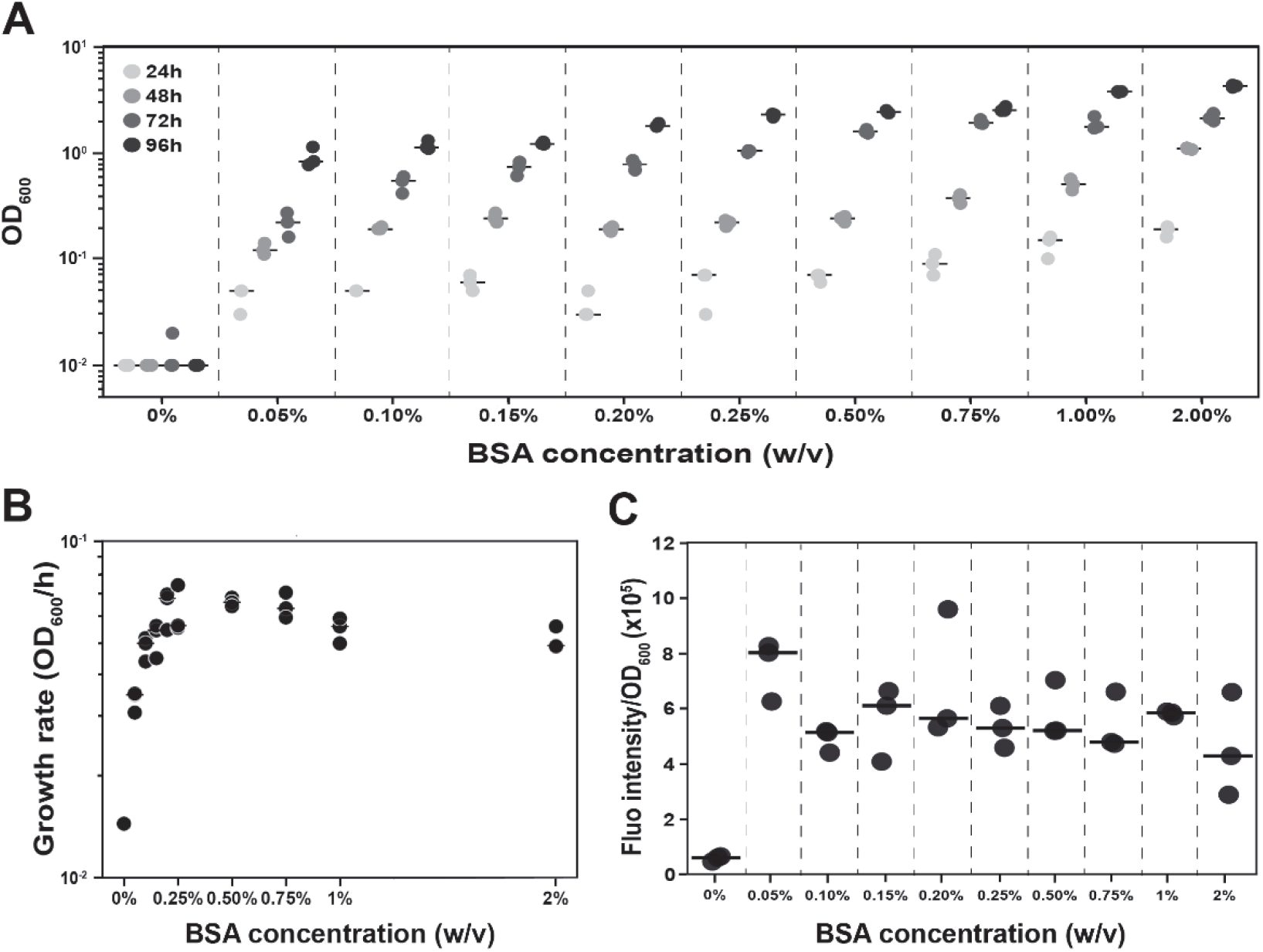
BSA concentration influence the yield of growth but not the EPs activity. **A**. Yield of the NCIB 3610 strain obtained at different time points in media containing 0.5% glycerol (v/v) with a BSA concentration ranging between 0% to 2% (w/v). Points represent OD_600_ values (n=3) and lines represent the median. **B**. The growth rate obtained from exponential regression performed on the yield of growth values displayed in (A). Each point represents the growth rate calculated for each replicate (n=3) for all BSA concentration ranging between 0% to 2% (w/v). The lines represent the median. **C**. Extracellular protease activity in the supernatant normalised to the total yield for a BSA concentrations ranging between 0% to 2% (w/v). Points represent fluorescence intensity/OD_600_ (n=3), and lines represent the median.

### Assessing the contribution of individual proteases to growth

The analysis detailed above reveals the collective impact of deleting all eight genes encoding the extracellular proteases from the genome of NCIB 3610. We next explored the impact on growth when (i) each coding region was deleted individually (Table 1), and (ii) when each extracellular protease coding region was individually reintroduced into its native locus on the Δ8 genome (Table 1, Fig. S3A). We measured growth attained by each strain after 96 hours in the presence of glycerol and 1% (w/v) BSA. We observed that there was a limited impact of deleting any single extracellular protease encoding gene, with a modest, but consistent, reduction in growth yield at 96h observed (Fig. 4A). In contrast, individually returning the coding regions for *aprE, bpr, epr, mpr, vpr*, or *wprA* into the Δ8 strain allowed for a partial recovery of growth when using BSA as the sole nitrogen source (Fig. 4B). There was no recovery of growth when the *nprB* or *nprE* coding regions were reintroduced to the genome (Fig. 4B). The ability of the spent culture supernatant harvested from the mono-producer strains to digest BSA was tested; only the *mpr* single producer strain showed any visible, albeit partial, digestion of BSA (Fig S3B). Partial BSA degradation was absent after enzymatic inactivation by heat treatment (Fig. S3B). These data highlight that collective action of the extracellular proteases is required to fully support feeding of *B. subtilis* on polymeric materials.

**Figure 4.**
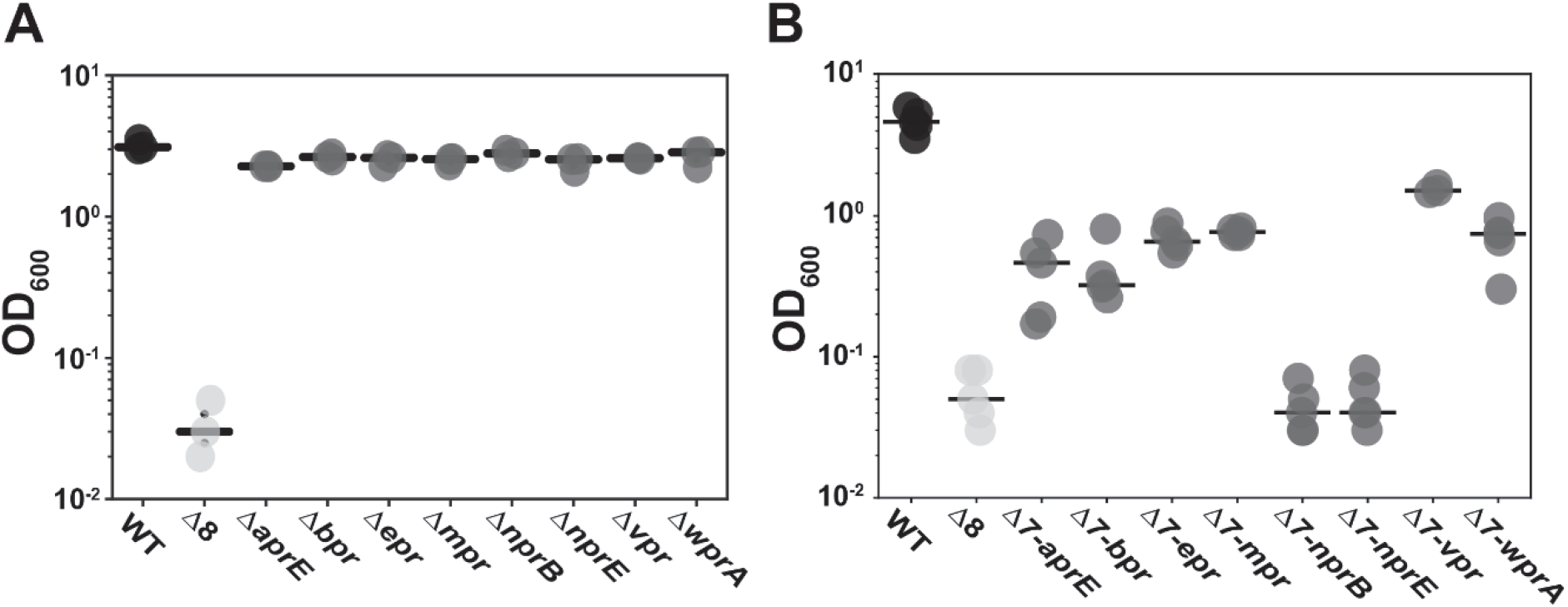
Collective role of extracellular proteases in supporting growth. Yield at 96h in media containing 0.5% (v/v) glycerol and 1% (w/v) BSA for NCIB 3610 (WT), NRS5645 (Δ8) and for **A**. single deletions strains *ΔaprE, Δbpr, Δepr, Δmpr, ΔnprB, ΔnprE, Δvpr, ΔwprA* (NRS3657, NRS6047, NRS3658, NRS3659, NRS3660, NRS3661, NRS7010, NRS6810 respectively) and for **B**. single extracellular protease producer strains *Δ7-aprE, Δ7-bpr, Δ7-epr, Δ7-mpr, Δ7-nprB, Δ7-nprE, Δ7-vpr, Δ7-wprA* (NRS3679, NRS3680, NRS3681, NRS 3882, NRS3683, NRS3684, NRS3685, NRS6362 respectively). In both cases each point represents the OD_600_ values (n=3 biological replicates and 1 technical replicate) and the lines represent the median.

### Extracellular proteases are a public good

We next tested if the extracellular proteases are a public good. We used a Transwell® system to physically separate WT and Δ8 strains within a shared, stationary growth environment. To allow quantification of growth in these conditions, we genetically modified the strains such that they constitutively produced either mKate2 or GFP (Table 1). We initially performed control experiments to assess that there was no cell diffusion between the Transwells® and the wells of the plates (Fig. S4A). Next, to ensure that the growth of *B. subtilis* was compatible with the Transwell® system, we used nutrient media conditions in which both strains could grow (glycerol and glutamic acid). We observed growth of the strains in both the upper and lower chambers of the Transwell®, irrespective of the strain inoculation combination used (Figure 5A, 5B and S4B, S4C).

**Figure 5.**
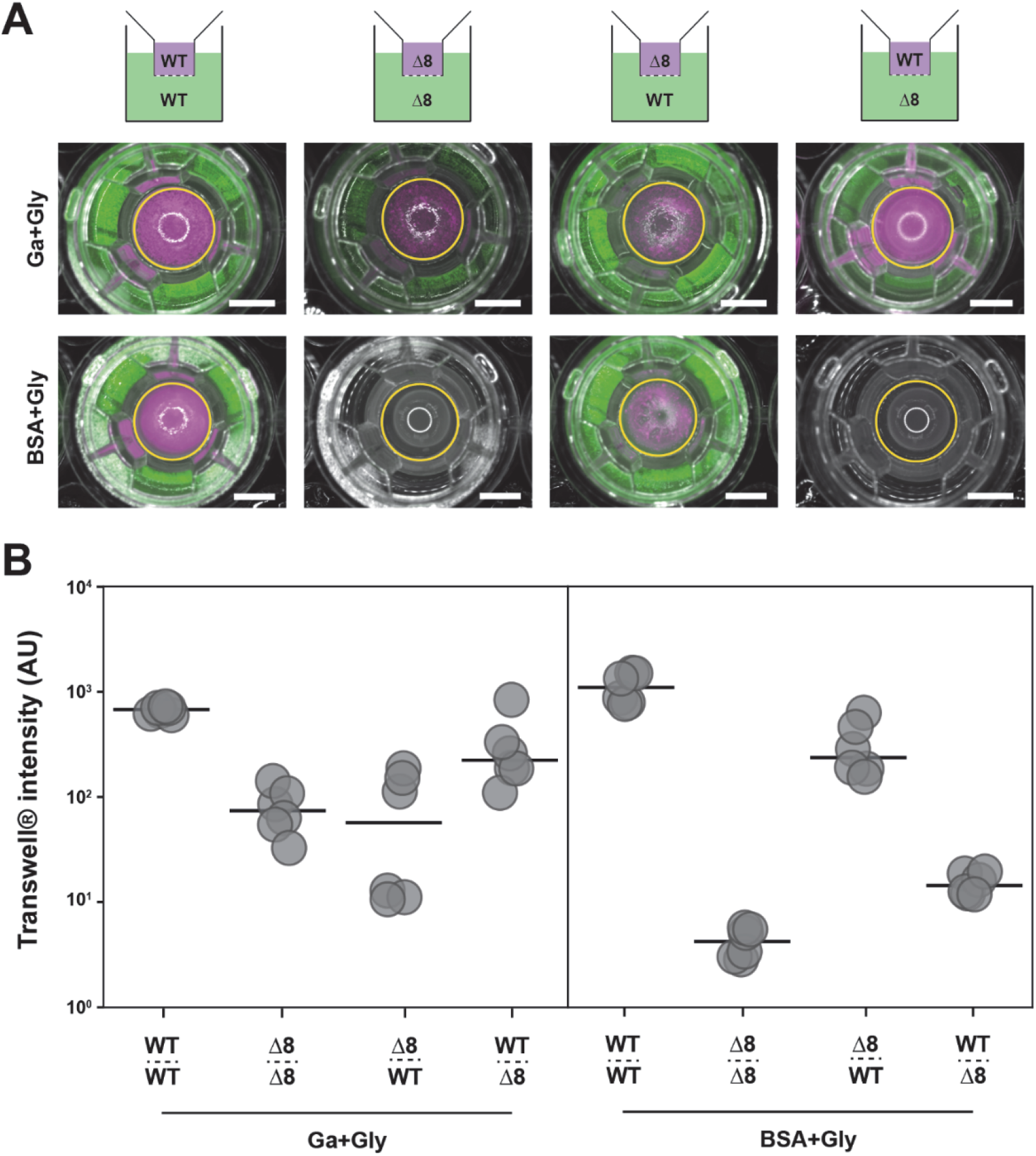
Extracellular proteases are a public good. **A**. A schematic of the experimental organisation. Transwell® assay merge images of reflected light, fluorescence from both the GFP (false coloured green) and mKate (false coloured magenta) channels after growth of the strain in 0.5% glutamic acid (w/v) and 0.5% glycerol (v/v) (Ga+Gly) (first row) and 1% BSA (w/v) and 0.5% glycerol (v/v) (BSA+Gly) (second row) media for different strain combinations. WT GFP (NRS1473) or WT mKate (NRS6932) and Δ8 GFP (NRS3648) or Δ8 mKate (NRS3670). The yellow circles represent the region of interest (ROI) used to quantify fluorescence values in the Transwell®. A representative image of three independent experiments is shown. The scale bar represents 5 mm. **B**. Transwell® fluorescence intensity after growth as detailed in **A**.. Conditions are annotated following this pattern: strain above dashed-line = Transwell® population, name below dashed-line = large well population. Each point represents fluorescence intensity values (n=3 biological replicates with 2 technical replicates) and the line represents the median value.

When the glutamic acid in the growth medium was substituted for BSA, as expected, the WT-WT strain pairings grew efficiently and no growth was observed for the Δ8-Δ8 strain pairings (Figure 5A, 5B and S4B, S4C). For the Δ8-WT strain combination, we observed robust growth of both strains when the Δ8 strain was within the smaller Transwell® and the WT in the large outer well (Figure 5A, 5B and S4B, S4C).

However, when the Δ8 strain was within the large outer well and the WT was in the small upper Transwell® no growth was measured for either strain (Figure 5A, 5B and S4B, S4C). These results allow us to make two conclusions: (i) extracellular proteases can act *distally* and therefore can be considered as a *public good*; the extracellular protease-producing NCIB 3610 strain can facilitate the growth of the extracellular protease-non-producing strain at a distance, and (ii) the co-culture can experience a *public good dilemma*; when the WT was limited to the smaller upper well, neither strain could grow indicating this configuration initiates an unsustainable balance of producer and non-producer to the ultimate detriment of both strains.

### The extracellular protease public good dilemma is context-dependent

We used a mathematical framework to further explore the occurrence of a public good dilemma in the context of extracellular protease production by *B. subtilis*. We devised a continuum framework comprising ordinary differential equations that describe the growth of wild-type cells *W*(*t*) and Δ8 (“cheater”) cells *C*(*t*) over time *t* ≤ 0. We assumed growth was within a well-mixed environment. Cells were assumed to grow (no extracellular proteases required) in response to an “available nutrient” *A*(*t*) that represented glutamic acid. It was assumed that BSA, represented by *B*(*t*), could not be directly consumed by cells. However, *B*(*t*) was assumed to be degraded by extracellular proteases, *E*(*t*) into a degraded form of BSA represented by *B*_*d*_(*t*). It was assumed that cells could grow in response to *B*_*d*_(*t*), but with a nutrient-to-biomass conversion rate less than that associated with glutamic acid *A*(*t*) (Fig. 6A). We tested the model for single-strain cultures across different simulated nutrient conditions (namely, those representing (i) glutamic acid and glycerol, (ii) glutamic acid and BSA, and (iii) BSA and glycerol) through appropriate choice of initial nutrient abundances. We found strong agreement between the growth of single-strain cultures *in-silico* and in experimental assays across all three conditions (compare Fig. S5 and Fig. 2).

**Figure 6.**
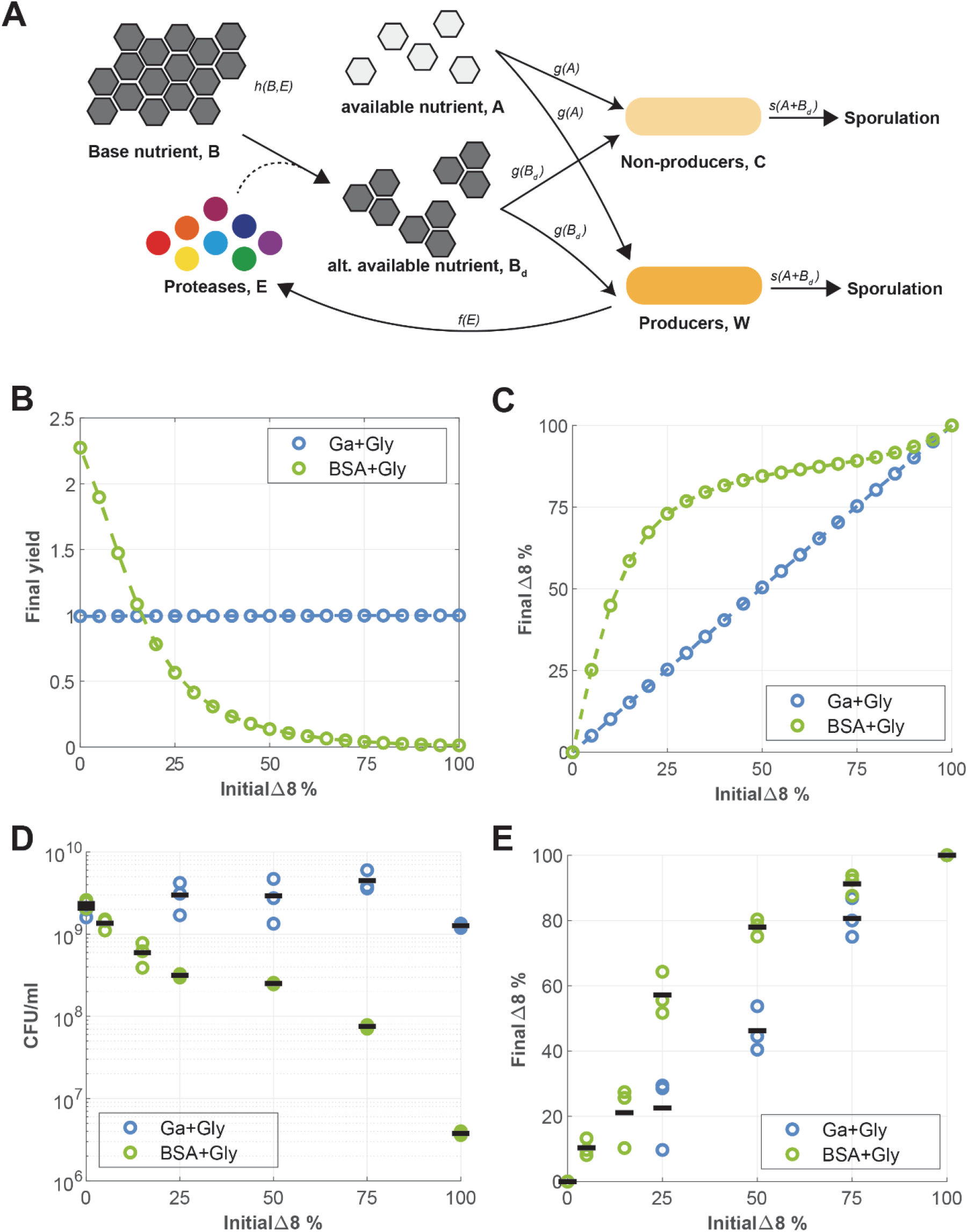
Exploration of the public good dilemma when extracellular proteases are shared. **A**. Schematic of the mathematical model. Producers, *W*, of extracellular proteases *E*, and non-producers *C*, can grow in response to two different nutrient sources; a nutrient *A*, representing glutamic acid, and an alternative nutrient *B*_*d*_, representing degraded BSA. Both producers and non-producers are assumed to sporulate in response to a lack of nutrient. Glutamic acid is represented by A in the model. BSA is represented by B. The nutrient A can be used directly by W and C. However, B requires the action of the protease E to convert it to B_and before it can be used by W or C. **B**. *In-silico* total yield (W+C+W_s + C_s) at *t* = 100 for different initial strain proportions. **C**. *In-silico* cheater proportion (% C+Cs) at *t* = 100 for changing initial strain proportions. **D**. CFU/mL representing the total population of WT-GFP (NRS1473) and Δ8-BFP (NRS3656) coculture in glutamic acid and glycerol (GA + Gly) (blue) and BSA and glycerol (BSA+Gly) (green) for different initial ratio of WT:Δ8 inoculum ranging from 0:100 to 100:0. Each point represents the total CFU/mL (n=3) and the lines represent the median. **E**. Final proportion of Δ8-BFP (NRS3656) population compared to initial Δ8-BFP (NRS3656) population from WT-GFP (NRS1473) and Δ8-BFP (NRS3656) coculture in glutamic acid and glycerol (GA + Gly) (blue) and BSA and glycerol (BSA+Gly) (green) for different initial ratio of WT: Δ8 inoculum ranging from 0:100 to 100:0. Each point represents the total CFU/mL (n=3), and the lines represent the median.

Next, we employed the model to investigate the potential outcomes of co-culturing the WT and Δ8 strains over a wide range of initial strain ratios. We defined the *in-silico yield* to be the *total* biomass density predicted by the model and includes both vegetative cells and spores as measured at the end of each simulation. For *in-silico* growth conditions representing the medium containing non-polymeric nutrient sources (glutamic acid and glycerol), the model predicted no change to the total yield as the initial proportion of Δ8 varied (Fig. 6B). However, for *in-silico* growth conditions representing the medium containing polymeric nutrients (BSA and glycerol), we observed a significant impact on total yield when changing the initial proportion of Δ8 (Fig. 6B). For example, the introduction of 10% Δ8 cells into a WT population led to a 35% decrease in total yield compared to a single-strain WT culture. Moreover, for an initial Δ8 ratio of 50%, total yield is reduced by a factor of 10, and for ratios above ∼75%, the total yield essentially saturates to zero. Hence, proportions of WT less than ∼25% are predicted to be incapable of supporting a dominant cheating strain with resultant collapse of both strains (Fig 5).

We also used our mathematical model to predict the relative abundance of both extracellular protease producers (WT) and non-producers (Δ8). The model predicts that if the extracellular proteases were not required for growth (glutamic acid and glycerol), the relative population proportions remained constant (Fig. 6C). By contrast, in the case where extracellular proteases were required for growth (BSA and glycerol), the relative abundance of Δ8 increased, no matter how large its initial proportion (Fig. 6C). Moreover, in line with the predicted effect on yield, larger impacts on final strain proportion were induced by the introduction of smaller initial proportions of Δ8 cells. Indeed, the greatest impact on final proportion was predicted to occur when the initial population contained ∼25% Δ8 strain; initial ratios around this value resulted in the greatest growth in Δ8 population. Hence the model reveals that the initial population ratio at which the cheater benefits the most matches the ratio that induces a significant reduction in total yield (down by ∼75%), but still just short of that value which would induce total population collapse (Fig. 6C).

Motivated by the model predictions, we experimentally analysed bacterial cultures grown with different WT to Δ8 starting ratios. Confirming our *in-silico* results, the findings show that sharing extracellular proteases with non-producing cells induced a reduction in total yield only when the extracellular proteases are needed to support growth (Fig 6D). Moreover, this reduction effect was found to respond in a non-linear manner to increasing the initial fraction of Δ8, with broadly the same saturating profile as that predicted by the model (cf. Figs. 6B and 6D noting the log scale in Fig. 6D). Inspection of the proportion of the two strains in the final culture revealed that if extracellular proteases are not required for growth, the relative population proportions remained constant (Fig. 6E). However, in the case that extracellular proteases are required for growth, the introduction of an initially small proportion of the Δ8 strain led to an increased final Δ8 strain fraction (Fig. 6E). Again, the “sweet spot” from the Δ8 perspective occurred at the 25% initial ratio data point. Together, these results suggest that the Δ8 strain has a fitness advantage over the WT due to the latter’s energy investment into extracellular proteases production. However, the impact of this growth advantage to the Δ8 strain is evidently dependent on the growth medium. Moreover, even when this growth advantage is apparent, its effect on the total yield is non-linear across the range of initial strain ratios and is vanishing when the fraction of the initial cheating population is either very low or close to 1.

The mechanism driving this complex response is elucidated in the next section.

### The relative cost of extracellular protease production underpins the public good dilemma

We have revealed the existence of a public good dilemma associated with the production of extracellular proteases that manifests only when these proteases are required for growth. This presents as a contradiction, because irrespective of the media choice, the WT population maintains similar levels of extracellular protease production when normalised to yield (Fig. 7A and B), presumably with the same metabolic (*absolute*) cost per extracellular protease unit. Therefore, we hypothesised that the selective penalty on growth incurred by the WT strain may be a result of the *relative cost* of extracellular protease production i.e., the *ratio* of the growth penalty associated with extracellular protease production relative to total growth. We hypothesised that this relative cost may be different in each of the media contexts.

**Figure 7.**
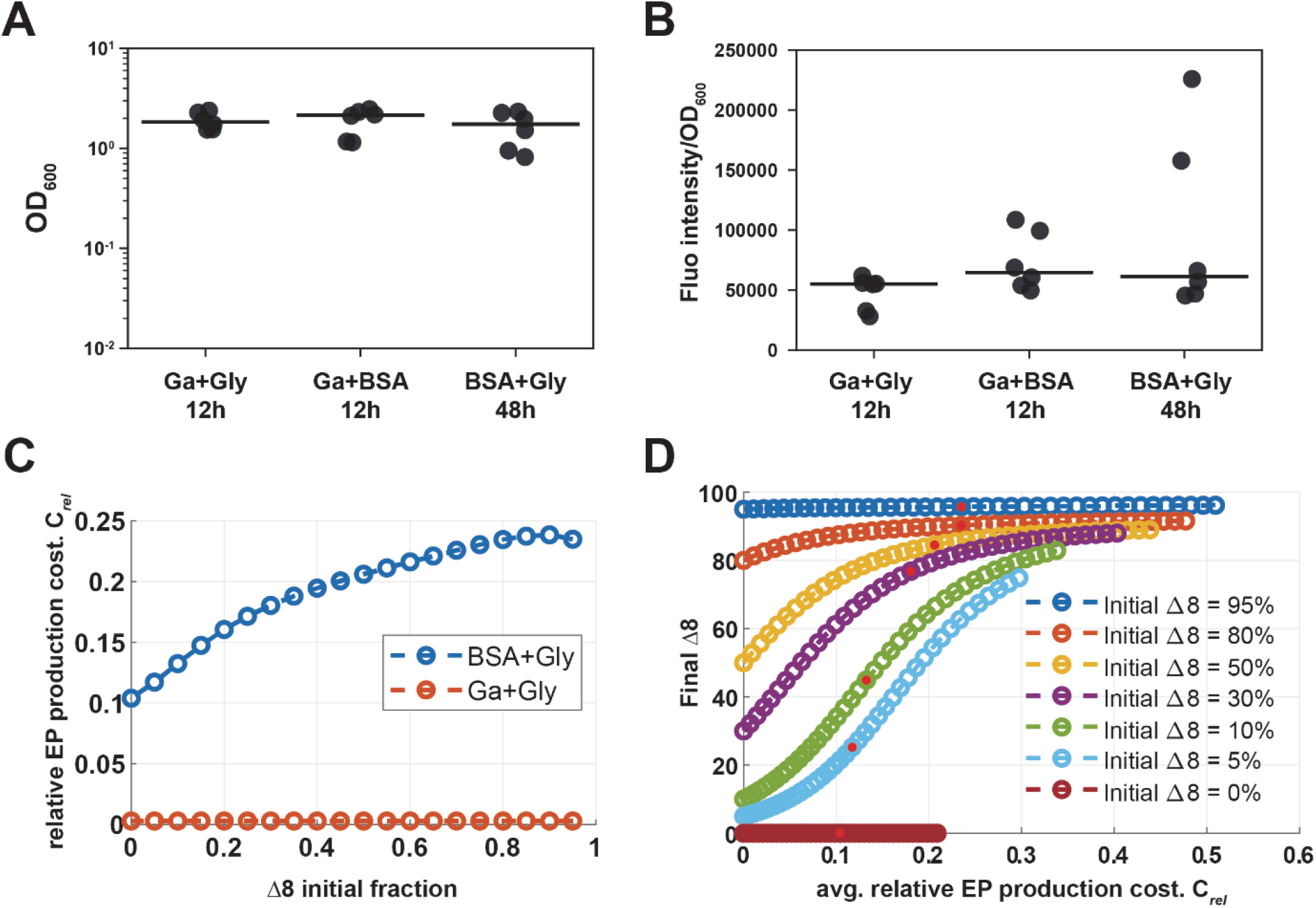
Relative cost of extracellular protease production explains the public good dilemma. **A**. Growth of NCIB 3610 in planktonic culture at 12h for glutamic acid 0.5% (w/v) and 0.5% glycerol (v/v) (Ga+Gly), 12h for 0.5% glutamic acid (w/v) and 1% BSA (w/v) (Ga+BSA) and 48h for 1% BSA (w/v) and 0.5% glycerol (v/v) (BSA+Gly). Points represent OD_600_ values (n=3 biological replicates with 2 technical replicates), lines represent median. **B**. Extracellular protease activity in the supernatant of culture from (A) normalised to the total yield represented in (A). Points represent fluorescence intensity/OD_600_ (n=3 biological replicates with 2 technical replicates), and lines represent the median. **C**. Relative cost of extracellular protease production against the initial fraction of Δ8 growth media representing glutamic acid and BSA as the nitrogen sources. **D**. *In-silico* proportion of the cheater at *t* = 100 against the *in-silico* relative cost of extracellular protease production during growth in a medium containing BSA as the sole source of nitrogen. Note that both axes represent model outputs. Data points correspond are generated by varying the value of *χ* (0 ≤ *χ* ≤ 2). Red dots represent *χ*_*s*_ = 1, the parameter value used in other simulations (see Table 4).

We tested this hypothesis by solving the model and comparing the computed *relative cost* in both media contexts across a range of initial WT to Δ8 population ratios, initially for a fixed absolute cost (*χ*).

The model simulations revealed that in conditions where the growth medium contains non-polymeric nutrients (glutamic acid and glycerol), the relative cost of extracellular protease production is small (< 0.01) for all initial Δ8 proportions (Fig. 7C). Recall that the WT and Δ8 strains performed equally well, with the final ratios closely matching their corresponding initial values (Fig. 6C). Thus, the model confirms that for non-polymeric growth media, a low relative cost correlates with the WT and Δ8 growing equivalently, irrespective of the initial Δ8 proportion. By contrast, when simulated in conditions representing the use of polymeric nutrient sources, the relative cost of extracellular protease production was computed to be at least an order of magnitude higher, across all initial Δ8 proportions, and to increase with the Δ8 fraction (Fig. 7C). We tested the robustness of our conclusions by varying the absolute cost (parameter *χ*) from a hypothetical case of zero absolute cost (*χ* = 0), to twice the value used in all other simulations (*χ* = 2*χ*_*s*_). In simulated non-polymeric nutrient conditions, the relative cost remained small despite the range of values of the absolute cost. Additionally, the final ratio of the Δ8 strain remained essentially unaltered from its initial value (Fig. S6). By contrast, in the simulated polymeric nutrient source medium, increasing *χ* over the same range caused the relative cost to increase significantly for each initial Δ8 ratio (Fig. 7D). Moreover, as the relative cost for production increased, the final Δ8 fraction also increased from its initial value (Fig. 7D). For each initial ratio, the greatest increase in the final Δ8 proportion occurred at the highest relative cost value.

Combined, our simulations predict the public good dilemma to be a context (nutrient source) dependent phenomenon. We demonstrated that for a fixed value of the absolute cost associated with extracellular protease production, the relative cost in each simulated media differed by at least an order of magnitude: when using non-polymeric nutrient sources, the relative cost for extracellular protease production is small, whereas in media requiring extracellular proteases for growth the relative cost is at least an order of magnitude higher. It is this high *relative* cost that determines a growth advantage to the cheater in this context and the higher this relative cost, the greater that advantage.

## Discussion

Here, using a combination of molecular genetics, physiological assays, and mathematical modelling, we provide evidence supporting the long-held, but previously unsubstantiated conjecture that the extracellular proteases of *B. subtilis* are a public good that support growth *via* the degradation of exogenous proteins. We established that in environmental conditions in which cells require extracellular proteases to grow, more than one of the extracellular proteases in the suite produced by the bacterium is required. Production of any single extracellular protease is insufficient to attain growth levels comparable to that of the wild-type strain and removal of any single extracellular protease coding region can be predominantly counteracted by the production of the remaining suite of proteases. We also provide evidence that the extracellular proteases function as a public good. Finally, we revealed that when growth is dependent on the activity of extracellular proteases, the extracellular protease-producing cells incur a significant cost of sharing this resource and the total yield of the whole community is reduced because of a public good dilemma.

### Dependence on extracellular proteases for growth

*B. subtilis* produces a suite of eight extracellular proteases, many of which have been shown to have specific molecular targets and roles (23, 33-36). Additionally, the extracellular proteases had been proposed to have a role in supporting growth, based on their ability to digest polymeric molecules (37), and by analogy with the mode of action of other enzymes, including chitinase and alginate lyase (9, 38). However, for *B. subtilis*, while the involvement of the extracellular proteases in sustaining survival in deep starvation (oligotrophic) conditions has been shown (24), explicit evidence supporting the role of the extracellular proteases in feeding when nutrients are abundant, but in a polymeric form, has been lacking. The construction of an extracellular protease-producing deletion strain, in an otherwise prototrophic strain background, allowed us to gain new insights into the growth-supporting role of extracellular proteases. We revealed the role exerted by each of the extracellular proteases in isolation through the generation of a suite of monoproducer strains (24, 39). We identified that individual reintroduction of six of the eight known extracellular protease coding regions allowed some recovery of growth. Interestingly, the Mpr monoproducing strain showed partial digestion of BSA and a partial recovery of growth, this is despite the fact that Mpr has been shown to be activated upon cleavage by Bpr (40). These results suggest that extracellular proteases may be self-activated, or show limited activity, in specific contexts, although further investigation is needed to fully understand their mode and extent of activation.

We also explored how *B. subtilis* balances its ability to sporulate with its ability to use extracellular proteases to sustain growth when presented with an abundant, but polymeric nutrient source. We uncovered that when presented with such a nutrient source, part of the population immediately sporulates. However, part of the population remains active and produces extracellular proteases that release usable nutrients from the polymeric source. Consistent with heterogeneous expression of the extracellular protease genes in the population (41) we assume that the release of nutrients subsequently allows the spores to germinate and growth of the population to occur. It will be of interest to explore the dynamics of sporulation and extracellular protease production in more depth in conditions where propagation of the cells is dependent on the activity of the enzymes to establish the timing of each process with respect to feedback from the local environment.

### The extracellular protease public good dilemma is context-dependent

*B. subtilis* produces many public goods including specialised metabolites with antimicrobial activity (42) and structural components of the biofilm extracellular matrix (43-45). The cost incurred by the cells when producing public goods is variable and can be context-dependent (46-48). Moreover, the extent to which public goods can be shared within and between populations varies (49-51). Here we showcase the public good attributes of *B. subtilis* extracellular proteases, uncovering that either the activity of the extracellular proteases, or the nutrients released after their action, can be shared with physically separated, non-producing cells within the same growth environment. We also established that the “public good dilemma” is only triggered in certain nutrient conditions, namely those in which nutrients are in polymeric form and growth is therefore dependent on the extracellular proteases. In these conditions, the producer cells incur a significant penalty on growth. However, there is apparently no measurable penalty to the wild type strain when it produces similar a level of extracellular proteases in conditions when they are not required for growth; there was no difference in growth yield to that observed for the Δ8 strain. This contrasts with other public goods produced by *B. subtilis*, for example the biofilm exopolysaccharide (51) where a significant growth advantage is afforded to non-producing cells (52) irrespective of if it provides an advantage or not.

Mathematical modelling allowed us to deduce that it is the *relative* cost of extracellular protease production that underpins the context dependency of the public good dilemma. When extracellular proteases are not required to support growth, the relative cost associated with their production is negligible. However, the slow-down of growth when cells use polymeric substrates as the source of nutrients increased the relative cost to a level that significantly impacted the growth rate.

Thus, the total yield of both wild type and non-producing cells was reduced. It may be that the public good dilemma is amplified by the fact that at least some of the genes involved in extracellular protease production are heterogeneously expressed (41, 53), meaning that even the WT extracellular enzyme producing population will contain cheaters. Overall, our results highlight the importance of the nutrient landscape in triggering a public good dilemma (9, 47), a situation that could significantly affect the development of a bacterial community (54, 55).

### Outlook

*B. subtilis* is a soil-dwelling bacterium that has been shown to live on both decaying plant matter (56, 57) and on living plant roots (58). Therefore, it is highly likely that extracellular proteases support growth in the bacterium’s natural environment by accessing polymeric nutrients released during decay and growth and/or generally contained within the soil itself (21, 59). As *B. subtilis* has promising applications as a biofertilizer (20, 60), it is important to understand how *B. subtilis* can settle and survive in diverse environmental conditions. We believe our findings will provide a foundation on which to build an understanding of how *B. subtilis* can survive in environments that diverge significantly in terms of nutrient accessibility.

## Acknowledgements

Work in the NSW laboratory is funded by the Biotechnology and Biological Science Research Council (BBSRC) [BB/P001335/1, BB/R012415/1]. TR’s PhD scholarship is funded by the Wellcome Trust [102132/B/13/Z]. We are very grateful to Dr. Sofia Arnaouteli for construction of strain NRS3685 and Dr. James Abbott for helpful conversations about genome submission processes. Genome sequencing was provided by MicrobesNG (http://www.microbesng.com).

## Data Availability Statement

Computational code and experimental datasets have been deposited in the nrstanleywall GitHub repository (https://github.com/NSWlabDundee/) and archived by BioStudies (61) and Zenodo (62). Sequencing datasets have been deposited in the ENA portal (PRJEB59494).

## Supplementary Figures

**Figure S1.**
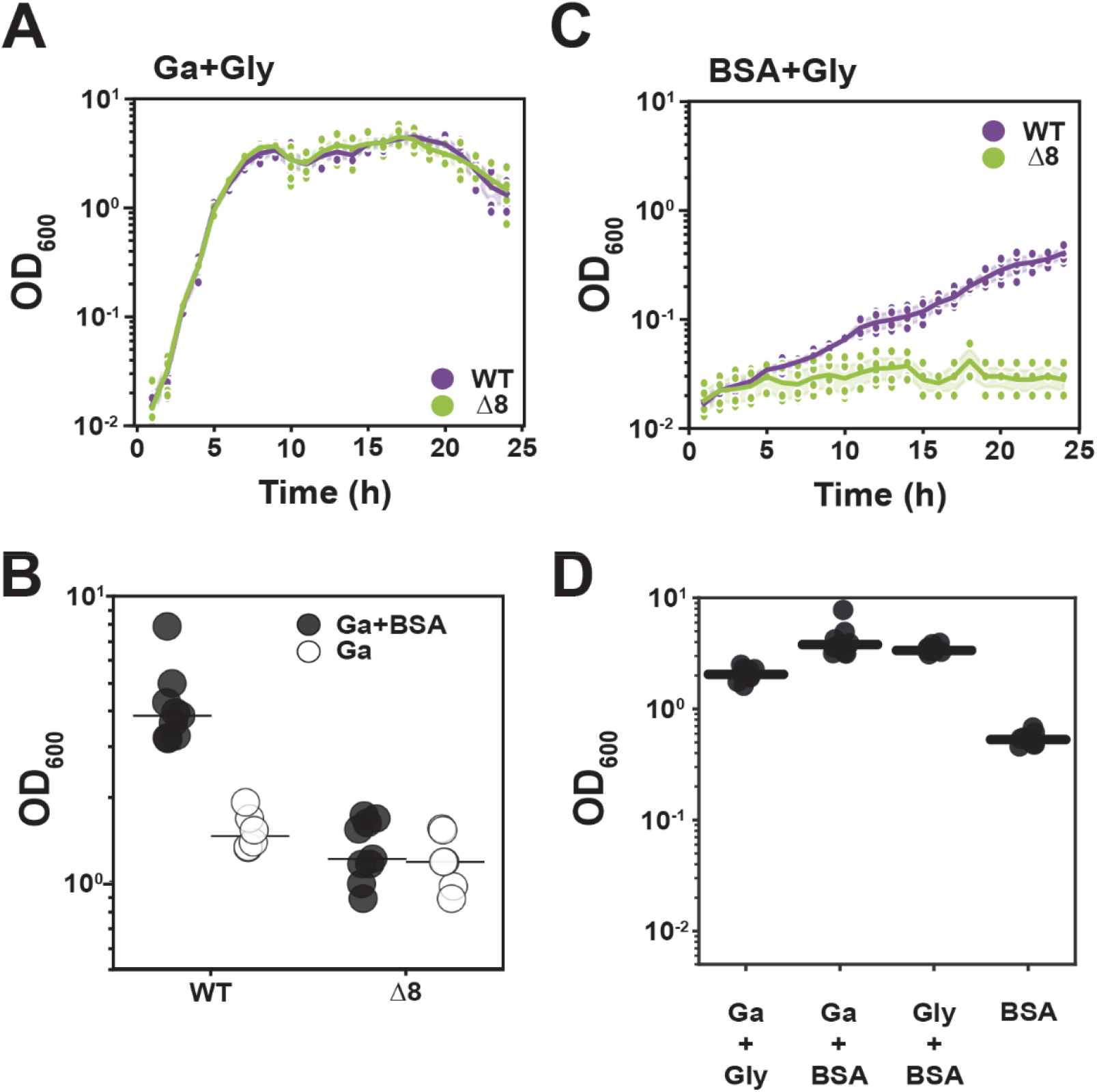
Growth analysis in different media. **A**. Growth curve of NCIB 3610 (WT) (purple) and NRS5645 (Δ8) (green) monoculture in 0.5% glutamic acid (w/v) and 0.5% glycerol (v/v) (Ga+Gly) (A) media between 0 and 24 h. Points represent OD_600_ values (n=3), lines represent the median and coloured-areas represent CI 95%. **B**. Yield of growth at 96h of NCIB 3610 (WT) and NRS5645 (Δ8) monoculture in 0.5% glutamic acid (w/v) and 1% BSA (w/v) (Ga+BSA) (black) and 0.5% glutamic acid (w/v) (Ga) (white). Points represent OD_600_ values (n=3 biological replicates with 2 technical replicates), and lines represent the median. **C**. Growth curve of NCIB 3610 (WT) (purple) and NRS5645 (Δ8) (green) monoculture in 1% BSA (w/v) and 0.5% glycerol (v/v) (BSA+Gly) (B) media between 0 and 24 h. Points represent OD_600_ values (n=3 biological replicates with 2 technical replicates), lines represent the median and coloured-areas represent CI 95%. **D**. Yield of growth at 96h for NCIB 3610 monoculture in 0.5% glutamic acid (w/v) and 0.5% glycerol (v/v) (Ga+Gly), 0.5% glutamic acid (w/v) and 1% BSA (w/v) (Ga+BSA), 1% BSA (w/v) and 0.5% glycerol (v/v) (BSA+Gly) and 1% BSA (w/v) (BSA) media. Each point represents the total CFU/mL (n=3 biological replicates with 2 technical replicates) and the lines represent the median.

**Figure S2.**
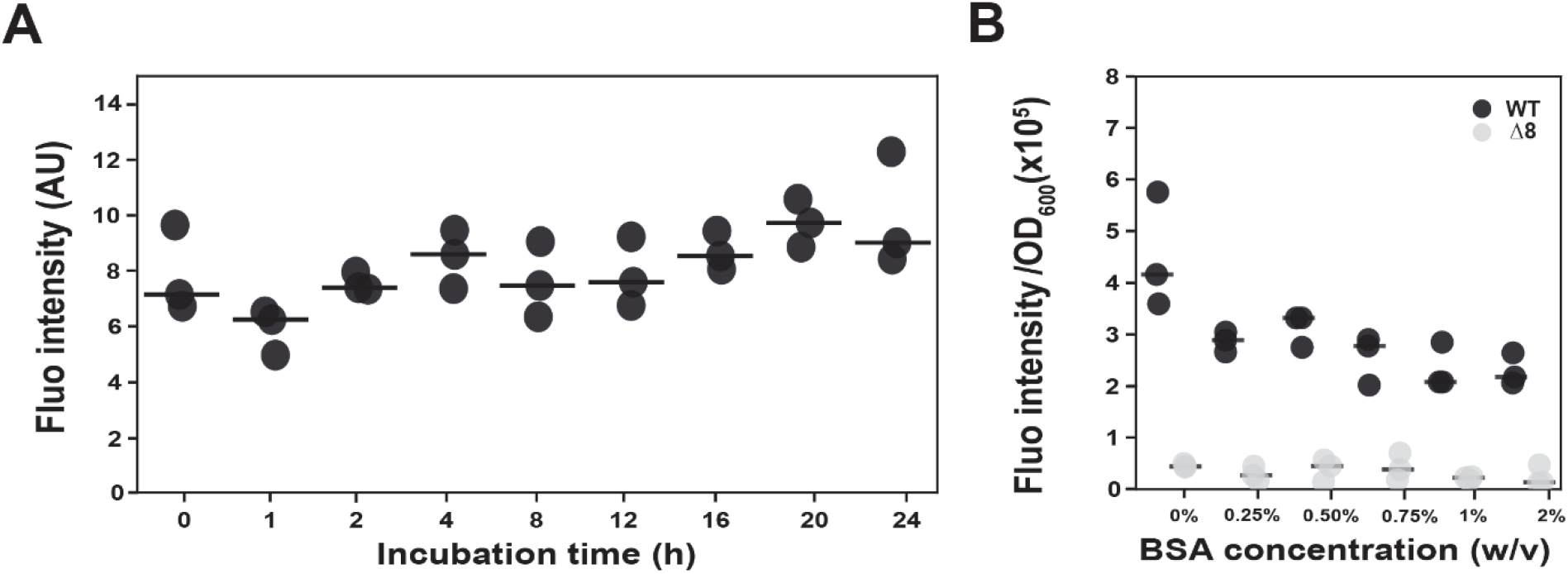
Extracellular protease activity assay controls. **A**. Extracellular proteases activity of culture supernatant obtained from NCIB 3610 monoculture in 0.5% glutamic acid (w/v) and 0.5% glycerol (v/v) (Ga+Gly) media at 24h and incubated at 37°C over a period time ranging from 0 to 24 h. Points represent fluorescence intensity (n=3) and lines represent the median. **B**. Extracellular proteases activity of culture supernatant obtained from NCIB 3610 monoculture in 0.5% glutamic acid (w/v) and 0.5% glycerol (v/v) (Ga+Gly) media at 24h in which BSA concentration ranging from 0 to 2% (w/v) was added during the protease activity quantification assay. Points represent fluorescence intensity (n=3) and lines represent median.

**Figure S3.**
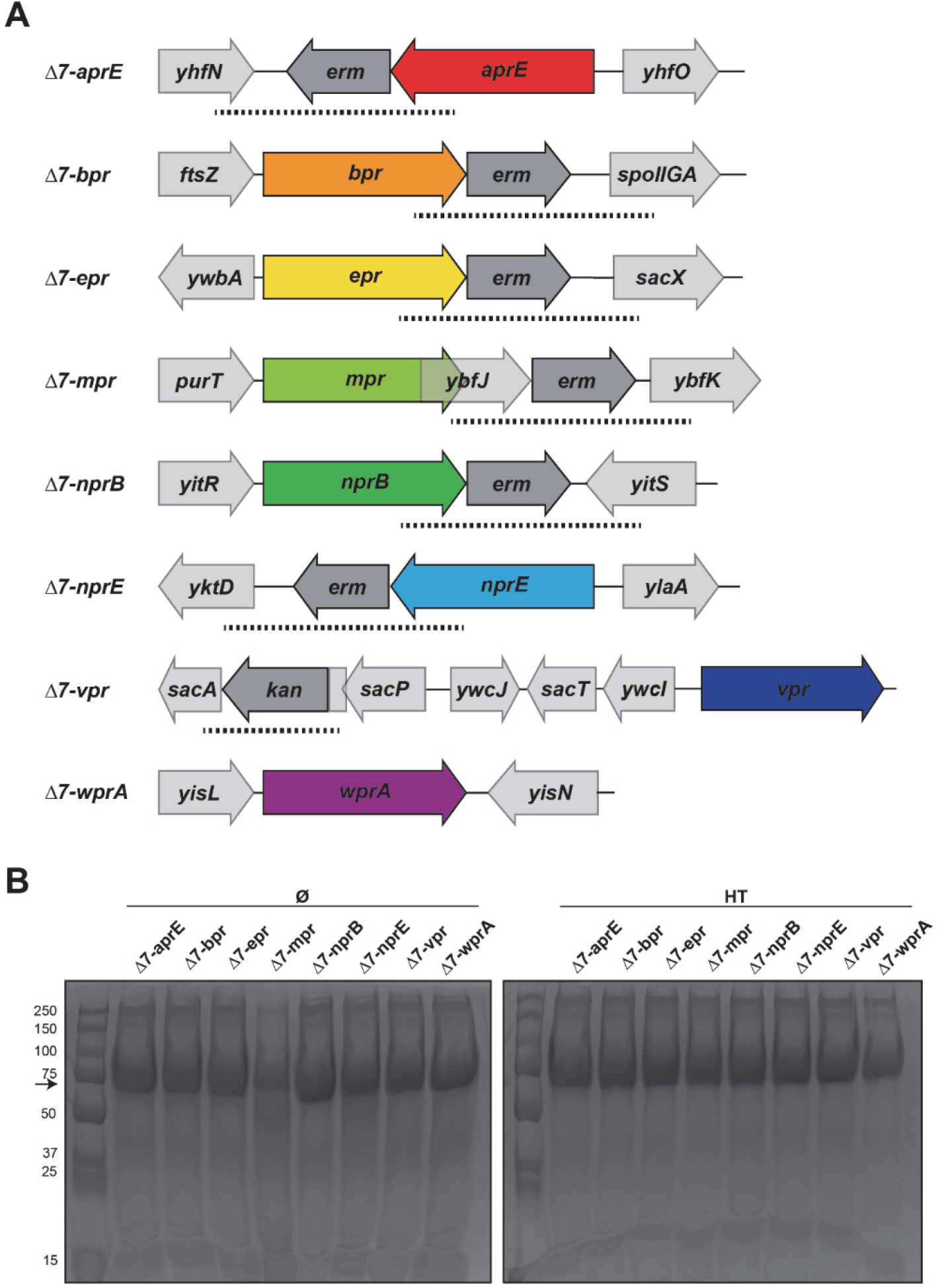
Monoproducer extracellular protease strains. **A**. Schematic of gene surrounding each extracellular protease interest for the monoproducer strains *Δ7-aprE, Δ7-bpr, Δ7-epr, Δ7-mpr, Δ7-nprB, Δ7-nprE, Δ7-vpr, Δ7-wprA* (NRS3579, NRS3680, NRS3681, NRS3682, NRS3883, NRS 3684, NRS3685, NRS6362, respectively). Dashed lines represent the region of DNA containing 500 bp of the extracellular protease gene region, erythromycin (*erm)* or kanamycin (*kan)* resistance gene and 500 bp of the downstream region of extracellular protease gene used for integration into the chromosome. **B**. BSA digestion assay using culture supernatant isolated after growth of the extracellular protease monoproducer strains *Δ7-aprE, Δ7-bpr, Δ7-epr, Δ7-mpr, Δ7-nprB, Δ7-nprE, Δ7-vpr, Δ7-wprA* (NRS3579, NRS3680, NRS3681, NRS3682, NRS3883, NRS 3684, NRS3685, NRS6362, respectively) in 0.5% glutamic acid (w/v) and 0.5% glycerol (v/v) (Ga+Gly) before (Ø) and after heat-treatment (HT). The black arrow represents BSA molecular weight (69 kDa). A representative image of three independent experiments is shown.

**Figure S4.**
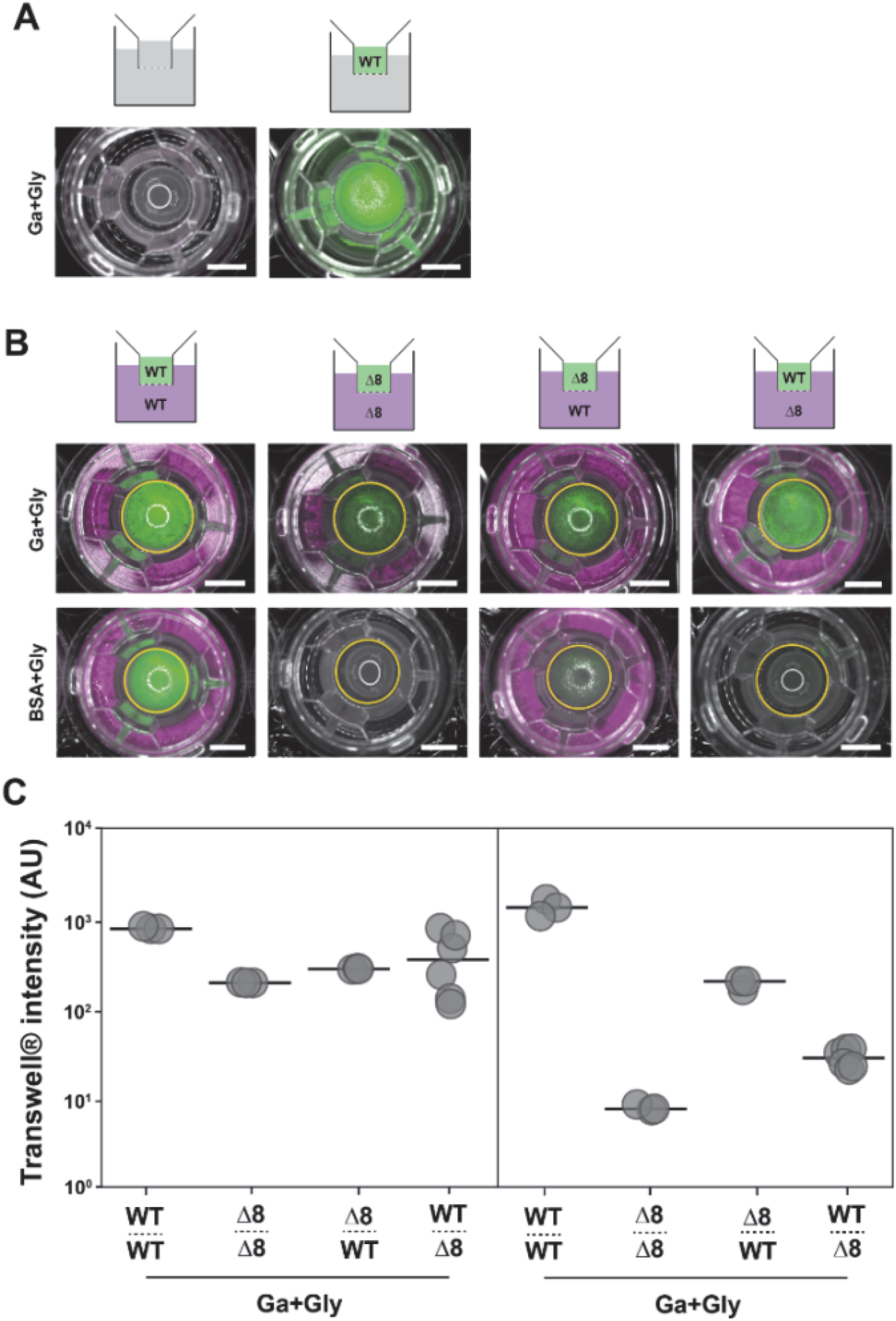
Reverse experiment of Transwell® assay. **A**. A schematic of the experimental organisation. Transwell® assay control merge images of reflected light and GFP (false coloured green) fluorescence channel after growth in 0.5% glutamic acid (w/v) and 0.5% glycerol (v/v) (Ga+Gly) media alone (right) and WT-GFP (NRS1473) (green) only inoculated in the Transwell® (left). A representative image of three independent experiments is shown. Scale bar represents 5 mm. **B**. A schematic of the experimental organisation. Transwell® assay merge images of reflected light, fluorescence from both the GFP (false coloured green) and mKate (false coloured magenta) channels after growth of the strain in 0.5% glutamic acid (w/v) and 0.5% glycerol (v/v) (Ga+Gly) (first row) and 1% BSA (w/v) and 0.5% glycerol (v/v) (BSA+Gly) (second row) media for different strain combinations. WT GFP (NRS1473) or WT mKate (NRS6932) and Δ8 GFP (NRS3648) or Δ8 mKate (NRS3670). The yellow circles represent the region of interest (ROI) used to quantify fluorescence values in the Transwell®. A representative image of three independent experiments is shown. The scale bar represents 5 mm. **C** Transwell® fluorescence intensity after growth in 1% glutamic acid (w/v) and 0.5% glycerol (v/v) (Ga+Gly) (left panel) and 1% BSA (w/v) and 0.5% glycerol (v/v) (BSA+Gly) (right panel). Conditions are annotated following this pattern: strain above dashed-line = Transwell® population, name below dashed-line = large well population. Each point represents fluorescence intensity values (n=3 biological replicates with 2 technical replicates) and the line represents the median value.

**Figure S5.**
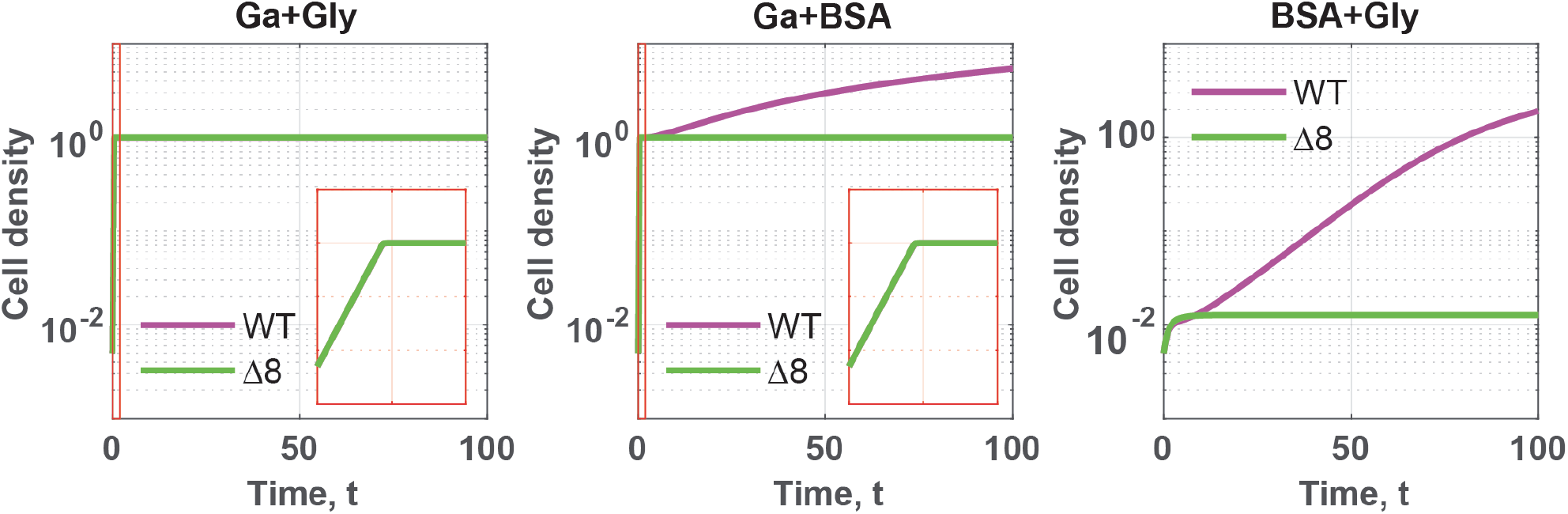
Supplemental *in-silico* data. Simulated single-strain cultures in different media. *In-silico* growth curves obtained from single-strain model simulations corresponding to the experimental data displayed in Fig. 2 A-C-E-G. The insets show a blow-up of the early growth dynamics. Parameter values are given in Table 4.

**Fig S6.**
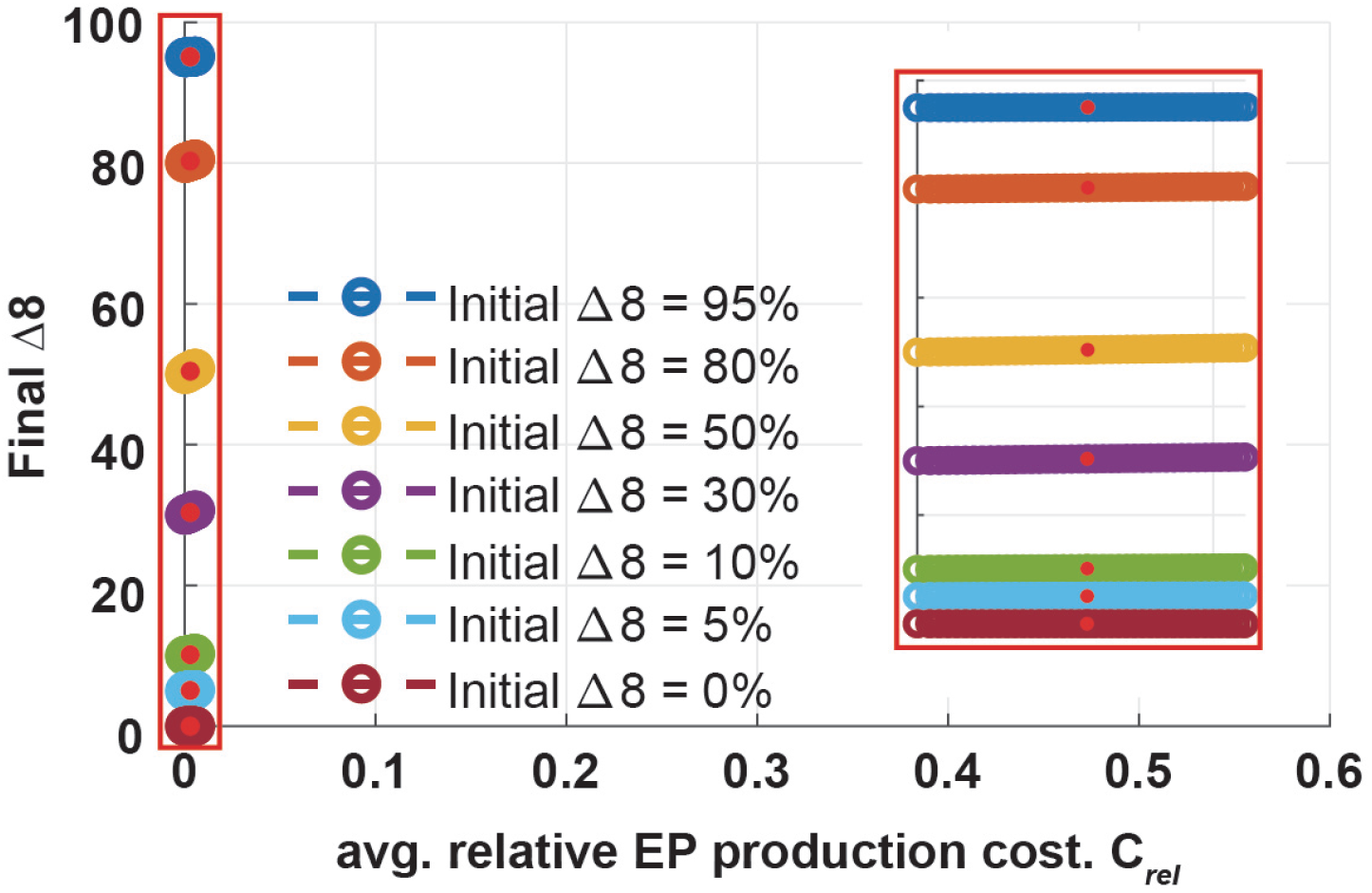
*In-silico* proportion of the cheater at *t* = 100 against the *in-silico* relative cost of extracellular protease production during growth in a medium containing glutamic acid as the sole source of nitrogen. Note that both axes represent model outputs. Data points correspond are generated by varying the value of *χ* (0 ≤ *χ* ≤ 2). Red dots represent *χ*_*s*_ = 1, the parameter value used in other simulations (see Table 4). Note that the x axis limits are identical to those used in Fig. 7C. The inset shows the dynamics for low relative costs.

## CRediT

| Term | Definition | Initials |
| --- | --- | --- |
| Conceptualization | Ideas; formulation or evolution of overarching research goals and aims | TR, LE, CE, FD, NSW |
| Methodology | Development or design of methodology; creation of models | TR, CE, LE, FD, NSW |
| Software | Programming, software development; designing computer programs; implementation of the computer code and supporting algorithms; testing of existing code components | TR, LE |
| Validation | Verification, whether as a part of the activity or separate, of the overall replication/ reproducibility of results/experiments and other research outputs | TR, CE, LE, FD, NSW |
| Formal analysis | Application of statistical, mathematical, computational, or other formal techniques to analyze or synthesize study data | TR, LE, FD, NSW |
| Investigation | Conducting a research and investigation process, specifically performing the experiments, or data/evidence collection | TR, LE |
| Resources | Provision of study materials, reagents, materials, patients, laboratory samples, animals, instrumentation, computing resources, or other analysis tools | TR, CE, FD, NSW |
| Data Curation | Management activities to annotate (produce metadata), scrub data and maintain research data (including software code, where it is necessary for interpreting the data itself) for initial use and later reuse | TR, LE, FD, NSW |
| Writing - Original Draft | Preparation, creation and/or presentation of the published work, specifically writing the initial draft (including substantive translation) | TR, LE, FD, NSW |
| Writing - Review & Editing | Preparation, creation and/or presentation of the published work by those from the original research group, specifically critical review, commentary or revision – including pre-or postpublication stages | TR, LE, CE, FD, NSW |
| Visualization | Preparation, creation and/or presentation of the published work, specifically visualization/ data presentation | TR, LE, FD, NSW |
| Supervision | Oversight and leadership responsibility for the research activity planning and execution, including mentorship external to the core team | LE, FD, NSW |
| Project administration | Management and coordination responsibility for the research activity planning and execution | FD, NSW |
| Funding acquisition | Acquisition of the financial support for the project leading to this publication | TR, FD, NSW |

